# IQD2 recruits KLCR1 to the membrane-microtubule nexus to promote cytoskeletal mechano-responsiveness in leaf epidermis pavement cells

**DOI:** 10.1101/2024.10.01.615909

**Authors:** Jonas Buhl, Sandra Klemm, Malte Kölling, Felix Ruhnow, Christian Ihling, Christian Tüting, Pradeep Dahiya, Jacqueline Patzsch, Leia Colin, Gina Stamm, Andrea Sinz, Panagiotos L. Kastritis, Staffan Persson, Katharina Bürstenbinder

## Abstract

Plant cells experience a variety of mechanical stresses from both internal and external sources, including turgor pressure, mechanical strains arising from heterogeneous growth between neighboring cells, and environmental factors like touch from soil, rain, or wind [1,2]. These stresses serve as signals at the cell-, tissue- and organismal level to coordinate plant growth during development and stress responses [3]. In plants, the physical cell wall-plasma membrane-microtubule continuum is proposed to be integral in transducing mechanical signals from the exterior to intracellular components [4–6]. Cortical microtubules (CMTs) rapidly reorient in response to mechanical stress to align with the maximal tensile stress direction [7,8]. Several studies proposed that CMTs themselves may act as stress sensors; the precise mechanisms involved in the regulation of CMTs and the modes of sensing, however, are still not clearly understood. Here, we show that IQD2 and KLCR1 are enriched at CMTs in proximity to the plasma membrane. IQD2, which is a *bona fide* microtubule-associated protein, promotes microtubule localization of KLCR1. By combining cross-linking mass spectrometry (XL-MS) and computational modeling with structure-function studies, we present first experimental insights into the composition and structure of IQD2-KLCR1 complexes. Further, we demonstrate that the IQD2-KLCR1 module is a positive regulator of microtubule mechano-responses in pavement cells. Collectively, our work identifies the IQD2-KLCR1 module as novel regulator of mechanostress-mediated CMT reorientation and provides a framework for future mechanistic studies aimed at a functional dissection of mechanotransduction at the plasma membrane-CMT interface during growth and plant morphogenesis.

**Highlights:**

- IQD2 and KLCR1 localize to the plasma membrane-microtubule nexus
- IQD2 is required for efficient microtubule targeting of KLCR1 *in planta*
- IQD2 physically interacts with KLCR1 and microtubules
- The IQD2-KLCR1 module promotes mechano-stress induced microtubule reorganization

## Results and discussion

Arabidopsis leaf epidermis pavement cells (PCs), which form highly complex jigsaw-puzzled shapes with undulating anticlinal walls, have become a popular model system to study how mechanical stress defines plant growth [9,10]. In PCs, cortical microtubules (CMTs) align with the maximal tensile stress direction in indenting neck regions where they guide plasma membrane (PM)-localized CELLULOSE SYNTHASE COMPLEXES (CSCs) for oriented deposition of cellulose fibrils [11]. This guidance enables stress-induced reinforcement of cell walls through microtubule-dependent cellulose deposition [12,13]. CMTs are anchored at the PM, likely via activities of various functionally distinct proteins [8]. This physical interaction may provide a basis for CMTs to sense and respond to mechanical stress, but experimental evidence supporting such functions is lacking. Previously, we identified two microtubule-localized proteins, IQ67 DOMAIN2 (IQD2) and KINESIN LIGHT CHAIN-RELATED1 (KLCR1), also known as CELLULOSE MICROTUBULE UNCOUPLING1 (CMU1), as novel regulators of PC shape [14,15]. IQD2 physically interacts with KLCR1 and the closely related KLCR2/CMU2 in yeast [14,16]. Since important and redundant roles of KLCR1 and KLCR2 in tethering of CMTs at the PM have been reported to stabilize CMTs against the pushing forces of CSCs [15,17], we tested if KLCR1 and KLCR2 also share functions during PC morphogenesis. To address this question, we analyzed PC shapes in cotyledons of wild type, *klcr1, klcr2* and *klcr1klcr2* mutants (Figure S1A-F) [18,19]. Consistent with our previous findings [14], PCs in *klcr1* mutants displayed increased circularity (Figure S1B,E) and increased largest empty circles (LEC) (Figure S1C,F) when compared to wild-type cells, which correlate with reduced shape complexity and increased mechanical stress, respectively, while cell area was not affected (Figure S1D). PCs in *klcr2* mutants were phenotypically indistinguishable from wild type, and *klcr1klcr2* double mutants resembled phenotypes of *klcr1* single mutants. These data demonstrate that KLCR1 functions independently of KLCR2 during PC morphogenesis.

### IQD2 and KLCR1 co-localize with microtubules at the cell cortex

The similarity in PC shape defects in *iqd2* and *klcr1* mutants and the indications for physical IQD2-KLCR1 interaction prompted us to explore their potential relationship at CMTs. To gain insights into the subcellular (co-)localization in Arabidopsis PCs, we examined the distribution of IQD2-GFP and KLCR1-GFP under control of the native promoters in the *ProIQD2:IQD2-GFP/iqd2* and *ProKLCR1:KLCR1-GFP/klcr1* complementation lines, which functionally rescue mutant phenotypes [14,20]. Using non-invasive confocal spinning disc fluorescence microscopy [21], IQD2-GFP and KLCR1-GFP fluorescence was observed in linear and punctate structures reminiscent of microtubules (Figure 1A,B,I,K). The mCherry (mCh)-TUA5 microtubule marker was introduced by crossing to analyze the distribution along the microtubule lattice [11]. IQD2-GFP and KLCR1-GFP were enriched in discrete puncta that clearly co-localized with mCh-TUA5, confirming microtubule localization of both proteins (Figure 1A,B,E,F). Co-localization was quantified by Pearson correlation coefficient (PCC) analysis, which showed that IQD2-GFP and KLCR1-GFP signals co-localized with microtubules (Figure 1C,G). Kymograph analyses revealed that IQD2-GFP and KLCR1-GFP foci were static, as indicated by vertical lines (Figure 1J,L). The immobile nature of IQD2-GFP and KLCR1-GFP-labeled foci was further supported by analysis of fluorescence recovery after photobleach (FRAP), which demonstrated reoccurrence of IQD2-GFP and KLCR1-GFP signals evenly along microtubules in punctate patterns (Figure S1G,H, Video V1 and V2). The dynamics of KLCR1 are in agreement with previous analyses of *ProUbiquitin10:GFP-KLCR1/CMU1* lines [15], demonstrating that KLCR1 localization is independent of the position of the GFP tag and the native vs *Ubiquitin10* promoter. Interestingly, we noticed that IQD2-GFP and KLCR1-GFP both were absent from microtubules detached from the PM at microtubule crossovers (Figure 1B,D,F,H, arrowheads). Based on our confocal micrograph analysis, we show that IQD2-GFP and KLCR1-GFP localize in static foci along microtubules at the cortex-PM nexus with similar spatial features.

**Figure 1:**
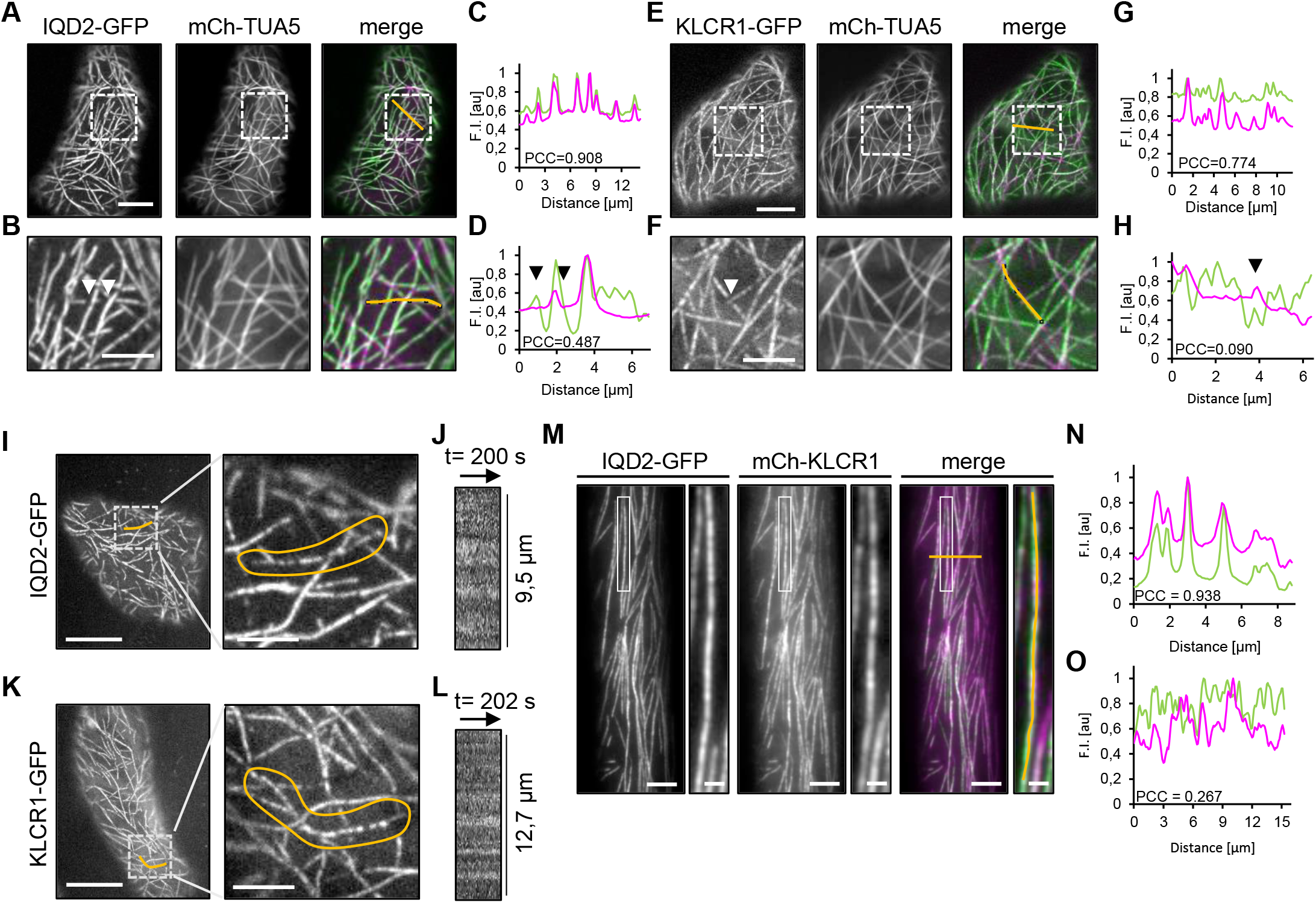
Subcellular localization and dynamics of IQD2 and KLCR1 at microtubules. **A**,**B**,**E**,**F**, Subcellular localization of IQD2-GFP (A,B) and KLCR1-GFP (E,F) relative to mCh-TUA5-labeled microtubules in the adaxial side of cotyledon epidermis cells of 7-day-old seedlings. Images are average projections (n = 100) of single z-planes (A,E) and magnification of framed regions in A,E (B,F). **C**,**D**,**G**,**H**, Pearson correlation coefficient (PCC) analysis of co-localization in fluorescence intensity profiles from mCh-TUA5 (magenta) with IQD2-GFP (green) (C,D) and KLCR1-GFP (green) (G,H) analyzed along the orange lines. **I-L**, Kymograph analysis of IQD2-GFP (I,J) and KLCR1-GFP (K,L) signals. Maximum projections of time series (left) and magnification of boxed regions (right) of IQD2-GFP (I) and KLCR1-GFP (K) in surface section of cotyledons epidermis cells of 10-day-old seedlings. Kymographs were generated from a line along the labeled microtubule of IQD2-GFP (J) and KLCR1-GFP (L). **M-O**, Co-localization of IQD2-GFP with mCh-KLCR1 in hypocotyl cell of 3-day-old seedlings by TIRF microscopy (M, left) and magnification of boxed area (M, right). Images are average projections (n = 100) Quantitative PCC analysis of co-localization in fluorescence intensity profiles from IQD2-GFP (green) and mCh-KLCR1 (magenta) along the orange line (N) and along a microtubule (O). Scale bars, 20 µm (I, left); 10 µm, (A,E); 5 µm (B,F,I (right),K,M); 1 µm, (M, magnification).

To test if KLCR1 and IQD2 foci spatially overlap along microtubules, we performed co-localization studies between IQD2-GFP and mCh-KLCR1 in crosses between the *ProKLCR1:mCh-KLCR1/klcr1* [17] and *ProIQD2:IQD2-GFP/iqd2* complementation lines. Analysis by spinning-disc microscopy revealed presence of mCh-KLCR1 and IQD2-GFP signals at microtubules in roots, hypocotyls, and cotyledon epidermis PCs (Figure S1I-L). For quantitative co-localization analysis at high resolution, we performed TIRF microscopy in hypocotyl cells, which are easier accessible for high-resolution imaging than leaf epidermis cells. Analysis of the outer periclinal cell cortex revealed partially overlapping localization of mCh-KLCR1 and IQD2-GFP in striate, punctate patterns (Figure 1M), for which we determined moderate PCCs for IQD2-GFP and mCh-KLCR1 fluorescence signals (Figure 1N,O), further pointing to a partial, yet incomplete overlap in subcellular localization. Thus, IQD2-GFP and mCh-KLCR1 exhibit comparable patterns and dynamics at the PM-CMT nexus and display partial co-localization indicative of both distinct and overlapping roles at microtubules.

### Microtubule-binding IQD2 recruits KLCR1 to the microtubule-PM nexus

The partial co-localization of IQD2 and KLCR1 at CMTs led us to further investigate the relationship between the two proteins. To test if subcellular localization of IQD2 and/or KLCR1 is dependent on their physical interaction in Arabidopsis, we introduced the IQD2-GFP and KLCR1-GFP constructs into the *iqd2klcr1* double mutant background. Analysis of IQD2-GFP localization revealed efficient microtubule targeting of IQD2-GFP in the *iqd2klcr1* background (Figure 2B), which was comparable to the *ProIQD2:IQD2-GFP/iqd2* complementation line (Figure 2A). By contrast, microtubule localization of KLCR1-GFP was strongly reduced in *iqd2klcr1* (Figure 2D) when compared to the *ProKLCR1:KLCR1-GFP/klcr1* complementation line (Figure 2C) and only few KLCR1-GFP-labeled puncta were detected at the cell cortex in *iqd2klcr1*. Quantitative analysis of KLCR1-GFP co-localization with mCh-TUA5 confirmed reduced microtubule targeting in *iqd2klcr1* (Figure 2E,F), as evidenced by PCC analysis (Figure 2G), and demonstrated accumulation of remaining KLCR1-GFP along microtubules (Figure 2F, insets). These findings indicate that IQD2 acts upstream of KLCR1 to facilitate KLCR1 microtubule tethering. In agreement with the partial but incomplete overlap of IQD2-GFP and mCh-KLCR1, these data further point to partially distinct KLCR1 functions at microtubules with and without IQD2. Interestingly, overexpression of RFP-KLCR1 under control of the strong CaMV 35S promoter in *Nicotiana benthamiana* leaves results in cytosolic accumulation of KLCR1, as reported previously [14,22,20], while co-expression with GFP-IQD2 induces a redistribution of RFP-KLCR1 to microtubules (Figure 3D). These data indicate that KLCR1 has no or only weak microtubule binding activity and that high levels of KLCR1 in the absence of sufficiently high amounts of IQD-linker proteins exceed the cell’s capacity to efficiently recruit KLCR1 to microtubules.

**Figure 2:**
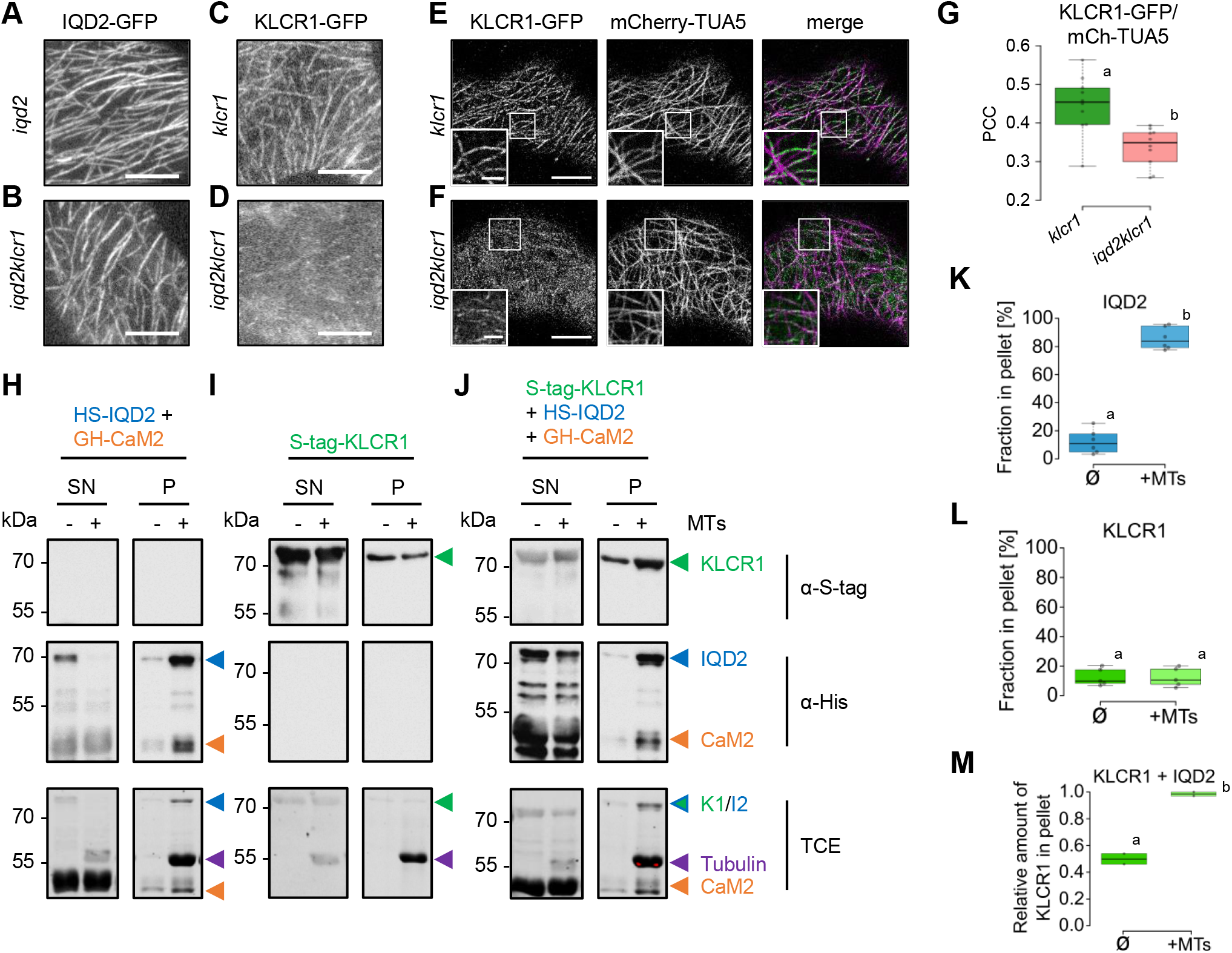
IQD2 is required for efficient microtubule recruitment of KLCR1. **A, B**, IQD2-GFP localization in cotyledon epidermis cells in the *ProIQD2:IQD2-GFP/iqd2* complementation line (A) and in the *ProIQD2:IQD2-GFP/iqd2klcr1* mutant (B) background. **C**,**D**, KLCR1-GFP localization in cotyledon epidermis cells in the *ProKLCR1:KLCR1-GFP/klcr1* complementation line (C) and in the *ProKLCR1:KLCR1-GFP/iqd2klcr1* mutant (D) background. **E-G**, Localization of KLCR1-GFP relative to mCherry-TUA5-labeled microtubules in the *ProKLCR1:KLCR1-GFP/Pro35S:mCh-TUA5/klcr1* (E) *and ProKLCR1:KLCR1-GFP/Pro35S:mChTUA5/iqd2klcr1* (F) background. Quantitative analysis of KLCR1-GFP colocalization with mCherry-TUA5 by PCC, n = 10 (G). Images are maximum projections of several z-planes (10 frames average each plane) Scale bars, 10 µm, A-D; 3 µm, Box in D. **H-M**, *In vitro* interactions of recombinantly expressed S-tag-KLCR1 (H), His-Sumo-IQD2 (HS-IQD2) coexpressed/copurified with GST-His-CaM2 (GH-CaM2) (I) and of S-tag-KLCR1 in the presence of HS-IQD2 and GH-CaM2 (J) with taxol-stabilized microtubules in spin down assay. Left, soluble fraction (supernatant); right, sedimented fraction (pellet) in the absence (-) and presence (+) of tubulins. Presence of S-tag-KLCR1 (top) and of HS-IQD2 and GH-CaM2 (center) was analyzed by immunodetection using anti-S-tag and anti-His antibodies, respectively. TCE-stained gels are shown on the bottom. Quantitative analysis of protein abundance of HS-IQD2 alone, n = 6 (K), of S-tag-KLCR1, n = 5 (l) and of S-tag-KLCR1 in combination with HS-IQD2/GH-CaM2, n = 2 (M) in pellet fractions in the absence (-) and presence (+) of microtubules. Different letters denote significant statistical differences by ANOVA with TukeyHSD (p<0.01).

**Figure 3:**
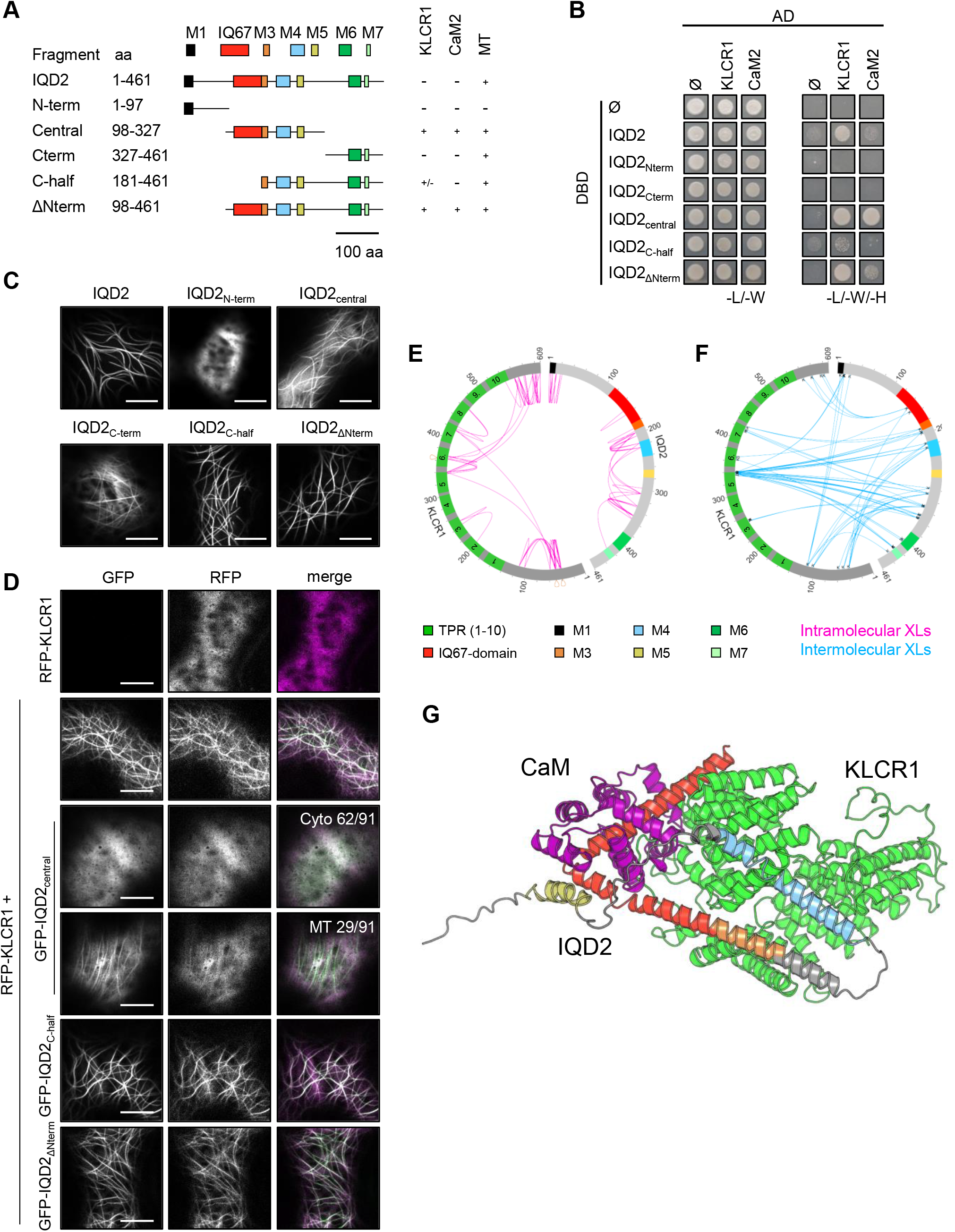
Structural determinants of IQD2-KLCR1 binding and microtubule targeting. **A**, Domain structure of IQD2 with motifs and domains indicated by colored boxes, and overview of truncated variants. + and – refer to the presence or absence of interactions, respectively, with KLCR1, CaM2, and microtubules (MT). **B**, Gal4-based yeast-two-hybrid assay with IQD2 variants fused to the Gal4 DNA-binding domain (DBD) and KLCR1 and CaM2 fused to the Gal4 activation domain (AD). –L/-W, vector-selective media; -L/-W-H, interaction-selective media. Yeast growth 5 days after spotting. **C**, Subcellular localization of GFP-fused IQD2 variants expressed under control of the CaMV 35S promoter in *N. benthamiana* leaves. **D**, Subcellular localization of *Pro35S:RFP-KLCR1* alone and upon coexpression with GFP-IQD2 variants. Numbers for GFP-IQD2_central_ fragments, co-expressing with RFP-KLCR1, represent the proportion of cells showing particular subcellular localization. Cyto, cytoplasm; MT, microtubules from n = 91 cells. Scale bars (C,D), 10 µm. **E**,**F**, Circular depiction of crosslink (XL) patterns in IQD2-KLCR1 complexes in the presence of CaM2. Intra-protein XLs along KLCR1 and IQD2 (E) and inter-protein XLs along IQD2-KLCR1. XLs were identified in at least 2/3 independent experiments. **F**, Superimposed structure of CaM2:IQD2:KLCR1 based on XL-MS-assisted docking and Alphafold prediction. Motifs and domains in IQD2 are colored according to the scheme in A; CaM2 is colored in magenta; KLCR1 in green.

Notably, recombinant His-tagged KLCR1 co-sediments with microtubules *in vitro* indicative of direct binding [15]. To test if physical interaction with IQD2 stabilizes KLCR1 at microtubules, we performed *in vitro* binding assays with taxol-stabilized microtubules [23]. KLCR1 was expressed in *Escherichia coli* with an N-terminal GST-His_6_-S-tag and purified using tandem-affinity tag purification, on-column GST-His_6_-tag-removal by thrombin cleavage, and subsequent size-exclusion chromatography (Figure S2A-D). His_8_-SUMO (HS)-IQD2 was co-expressed and co-purified with its binding partner GST-His_6_(GH)-Calmodulin2(CaM2) (Figure S2G,F), which greatly improved the yield and solubility of recombinant HS-IQD2 (Figure S2E,F), using a tandem affinity purification protocol. A commercially available MICROTUBULE-ASSOCIATED PROTEIN (MAP) fraction (MAPF) and BSA were included as positive and negative controls, respectively (Figure S2H). As expected, MAPF was enriched in pellets co-sedimenting with microtubules while BSA was independent of the presence or absence of microtubules. A small fraction of HS-IQD2 and S-tag-KLCR1 sedimented in the absence of microtubules (Figure 2H,I, bottom). The amount of HS-IQD2 in the pellet fraction, however, was significantly higher upon addition of polymerized tubulins (Figure 2H,K). GH-CaM2 co-sedimented with microtubules in the presence of HS-IQD2 (Figure 2H) while it remained soluble in the absence of HS-IQD2 (Figure S2I). These data demonstrate that CaM2 does not bind microtubules directly, which is in agreement with earlier reports [24–26], and confirm the presence of IQD-CaM2 complexes at microtubules [14,22]. The ratio of soluble to sedimented S-tag-KLCR1 was not affected by the presence or absence of microtubules (Figure 2I,L), indicating that S-tag-KLCR1 does possess very weak or no microtubule binding capacities, which is in contrast to the previously reported microtubule binding of His-tagged KLCR1 [15]. These differences may be caused by the N-terminal His-tag, which significantly increases the affinity of the MAP END-BINDING 1 (EB1) for microtubules, and possibly has similar effects on KLCR1-microtubule interactions [27], or could be due to protein aggregations of the KLCR that appear to exhibit weak microtubule affinity. Notably, N-terminal GST-His_6_-tagged CaM2 remained soluble (Figure S2I); the His-tag alone thus is not sufficient for microtubule binding. When combined with HS-IQD2/GH-CaM2, the amount of S-tag-KLCR1 increased significantly in the pellet fraction (Figure 2J,M), demonstrating IQD2-mediated microtubule targeting of KLCR1. Together with the reduced microtubule localization of KLCR1-GFP in the *iqd2klcr1* mutant background, these data indicate partially IQD2-dependent recruitment of KLCR1 to microtubules. Residual microtubule targeting of KLCR1 in the absence of IQD2 may occur either via weak direct microtubule binding or could be mediated by additional PC-expressed *IQD* family members [28–31] or other KLCR-interacting proteins, such as members of the KINESIN 4 [17] or NETWORKED [14] family.

### Structural determinants of IQD2’s KLCR1 binding and microtubule recruitment

To gain insights into the mechanisms involved in IQD2-mediated microtubule-recruitment of KLCR1, we performed structure-function studies with N- and C-terminally deleted IQD2 variants (Figure 3A). IQD2 comprises the central and eponymous IQ67 domain, required and sufficient for CaM binding [22], and several short, conserved motifs (M) that are interspersed by hypervariable intrinsically disordered regions (IDRs) [30,32–34] (Figure 3A, Figure S3A). To map binding regions, we performed GAL4-based yeast two-hybrid assays between IQD2 variants and KLCR1. As control, CaM2 fusions were included, which interact with IQD proteins via the IQ67 domain [22,35]. Analysis of yeast growth revealed preferential interaction of KLCR1 with the IQD2_central_ and IQD2_ΔNterm_ fragments that both harbor the IQ67 domain and conserved motifs M3, M4, and M5 (Figure 3B), while co-expression of KLCR1 with the N- and C-terminal IQD2 variants (IQD2_Nterm_, IQD2_Cterm_) failed to restore yeast growth. Deletion of the IQ67 domain in the IQD2_C-half_ variant fully abolished CaM2 binding, but not KLCR1 binding. Physical interaction with IQD2_central_, IQD2_C-half_, and IQD2_ΔNterm_ was validated in *in vitro* pull-down assays between Ni-NTA-immobilized HS-IQD2 fragments and recombinant GST-KLCR1 compared to the GST negative control (Figure S3C,D). Together, our data demonstrate that the central region of IQD2 is sufficient, and that the IQ67 domain is not required, for KLCR1 binding.

Analysis of GFP-fused IQD2 variant localization in transient expression assays in *N. benthamiana* revealed the presence of two distinct microtubule-binding domains (MBDs) in the IQD2_central_ and the IQD2_Cterm_ variant, respectively, that were sufficient for partial microtubule targeting when compared to full-length GFP-IQD2 or the IQD2_ΔNterm_ variant (Figure 3C). These MBDs share similarity with MBDs identified in IQD8, IQD9, IQD13 and IQD16 [20,36–38]. Notably, the central part harboring MTB1 also contains the KLCR1 binding region (Figure 3A,B). To test if this region can simultaneously bind KLCR1 and microtubules, we performed co-expression assays in *N. benthamiana* between GFP-fused IQD2 variants and RFP-KLCR1 expressed under control of the CaMV35S promoter (Figure 3D). Upon co-expression of the MTB1-containing IQD2_central_ variant with RFP-KLCR1, GFP-IQD2_central_ lost its microtubule localization and accumulated in the cytosol in 68 % of the cells. The GFP-IQD2_C-half_ and GFP-IQD2_ΔN-term_ variants, in which the motifs M3, M4, and M5 and the C-terminal MTB2 are combined, efficiently mediated KLCR1 recruitment in all analyzed cells (Figure 3D). These data suggest that KLCR1 binding can interfere with microtubule targeting of MTB1, while the C-terminal MTB2 mediates anchoring of IQD2-KLCR1 complexes at microtubules.

To explore the interaction between IQD2 and KLCR1 in more detail, we aimed to gain insights into the relative orientation and arrangement of the proteins. Primary sequence analysis suggests that IQD2 and KLCR1 both contain IDRs (Figure S3A,B,F-I). The N- and C-terminal disordered linker of KLCR1 presumably allows the protein to explore an extended conformational space [39,40]. The central region is predicted to adopt a canonical tetratricopeptide repeat (TPR) domain fold, consisting of a superhelical arrangement of helix-turn-helix repeats that mediate protein-protein-interactions and often function in assembly of macromolecular complexes (Figure S3B,G) [41,42]. As proteins rich in IDRs lack stable conformations and exist in ensembles of conformations that are difficult to assess in structural biology studies, we performed chemical XL-MS experiments [43,44]. After purification, IQD2/CaM2 and KLCR1 were chemically cross-linked with the MS-cleavable cross-linker disuccinimidyl dibutyric urea (DSBU) that connects protein amine groups (lysine residues and N-termini) with Cα-Cα distances of 30 Å [43]. Coomassie staining of protein samples indicated efficient cross-linking by DSBU as demonstrated by the accumulation of higher molecular weight species corresponding to IQD2:CaM2 and KLCR1:IQD2:CaM2 complexes (Figure S3E). SDS gel bands corresponding to the assembled KLCR1:IQD2:CaM2 complexes were excised and subjected to LC/MS analysis to identify cross-linked residues in KLCR1, IQD2, and the KLCR1:IQD2:CaM2 complex. In KLCR1, we uncovered short-range cross-links within the putative IDRs in the N- and C-termini and several cross-links within single and between neighboring TPR motifs. This finding is consistent with the predicted α-helical structure and proximity of adjacent TPR motifs in TPR domains (Figure 3E) [42]. In IQD2, short range cross-links were identified between adjacent motifs and flanking IDRs, e.g., between M1-flanking IDR, and M4-flanking IDRs (Figure 3E). Additional cross-links were found between IQD2 with the region 338 to 364 of KLCR1, corresponding to TPR5 and TPR6. Interestingly, the C-terminal IDR of IQD2 was part of the interaction between these two proteins (Figure 3F). On top of that, several cross-links involving the IDRs of both proteins indicate a stabilization and structural compaction within the protein complex. Although CaM2 was present in the cross-linking reaction, no cross-links involving CaM2 were discovered. It has to be noted that absence of cross-links does not exclude complex formation.

To generate atomic models of the KLCR1:IQD2:CaM2 complex, AlphaFold2 models were used as input (Figure S3F-I) [45] and molecular docking was performed with the HADDOCK webserver [46]. To prepare the models, the disordered linkers, displayed with low confidence (plDDT and PAE-score) were removed prior docking. For IQD2 and KLCR1, the residue ranges of 114 to 291 and of 30 to 606, respectively, were considered. Cross-linking data were sufficient to determine two distinct clusters. Upon calcium binding, holo-CaM can bind target peptides in which the linker helix undergoes a hinge-like motion and folds around the target sequence. Based on published models, CaM can bind to helical elements as well as helix-turn-helix motifs [47]. AlphaFold2 prediction of an IQD2:CaM2 complex suggests the latter, that the previous extended helical element of IQD2 is actually in a helix-turn-helix motif (Figure 3G). This is also supported by a higher PAE score for this interaction region, adding confidence to the predicted structure (Figure S3J,K). Overlaying this structure with retrieved docking results reveals that the N-terminal region of IQD2 is positioned near KLCR1, which is corroborated by crosslinks between IQD2’s IDR to KLCR1 (Figure 3G). These particular crosslinks are incompatible with an extended α-helix as the N-terminal area of IQD2 is distant from KLCR1 in such a conformation. Overall, these data point to a direct, physical interaction between the central region of IQD2 with the central TPR motifs of KLCR1 that likely is stabilized by additional interactions mediated by the IDRs, while the C-terminal MTB2 of IQD2 is required to tether the IQD2-KLCR1 complex at microtubules.

### The IQD2-KLCR1 module promotes cytoskeletal reorganization in response to mechanical stress

The physical interaction and colocalization of IQD2 and KLCR1 at the CMT-PM nexus and the evidence for IQD2-mediated microtubule targeting of KLCR1 prompted us to assess the genetic interaction and the contribution to microtubule organization during PC morphogenesis in *iqd2klcr1* mutants. Consistent with previous reports, we observed defects in PC shape in *iqd2* and *klcr1* mutants compared to wild type [14], as reflected by an increase in circularity and LEC, respectively (Figure S4A-F). The phenotypic differences were not further enhanced in *iqd2klcr1* suggesting that IQD2 and KLCR1 act together in PC morphogenesis. To assess microtubule organization in PCs, we introduced the mCh-TUA5/LTi6b-GFP dual microtubule/PM marker into the *iqd2klcr1* background [48]. Analysis of microtubule organization in PCs (Figure 4A-F) revealed no significant differences in the mean values of microtubule anisotropy between wild type and *iqd2klcr1* in whole cells (Figure 4D) or within the LECs (Figure 4B,E). Similarly, means of normalized gray values, corresponding to microtubule signals, within regions adjacent to lobes and necks were comparable between wild type and the *iqd2klcr1* mutant (Figure 4C,F). In *iqd2klcr1*, however, microtubule anisotropy displayed significantly higher variance at necks, which correspond to highest stress regions along the undulating wall, while no differences were observed in lobes (Figure 4F). These differences are indicative of altered CMT organization in response to mechanical stress. We thus wondered whether the IQD2-KLCR1 module may be a general regulator of stress-induced microtubule organization and also affects CMT reorganization in response to tissue-scale mechanical stress imposed upon cell ablation [12,6]. As reported previously, CMTs in wild type responded by co-aligning with the border of the ablation site and by gradual increase in microtubule ordering (Figure 4G,H) [6]. In *iqd2klcr1*, microtubule reorganization was delayed at two and four hours after ablation, but reached wild-type levels six hours after ablation. These results indicate that the IQD2-KLCR1 module promotes microtubule reorganization in response to changes in mechanical stress, potentially by stabilizing CMT-PM contact sites. The mechanical coupling between CMTs and the PM by the IQD2-KLCR1 module, which possibly is further stabilized by lipid-anchoring of KLCR1 via S-acylation [49] likely primes CMTs for sensing or rapid responsiveness to mechanical stress, including the dynamic reorganization of microtubules and subsequent events in the signal transduction cascade. The functional significance of the IQD2-KLCR1 module presumably extends beyond PCs, as both proteins are ubiquitously present across all analyzed plant tissues (Figure S1I) [15,14]. Lastly, although the specific functions of calcium signaling in PCs remain unknown, calcium signals represent one of the most rapid responses to mechanical stress [50,51]. In animals, calcium is released from the endoplasmic reticulum (ER), which functions as an intracellular calcium store, and physical ER-PM contact sites (EPCS) play important roles in this process [52]. While the role of the ER and EPCS in plant calcium signaling is not yet clear, KLCR1 interacts with core components of plant EPCSs [14]. Examining the role of calcium-CaM in regulating IQD2-KLCR1 functions therefore presents an intriguing experimental platform to decipher the molecular mechanisms underlying the integration of calcium signals at CMTs during mechanotransduction [53].

**Figure 4:**
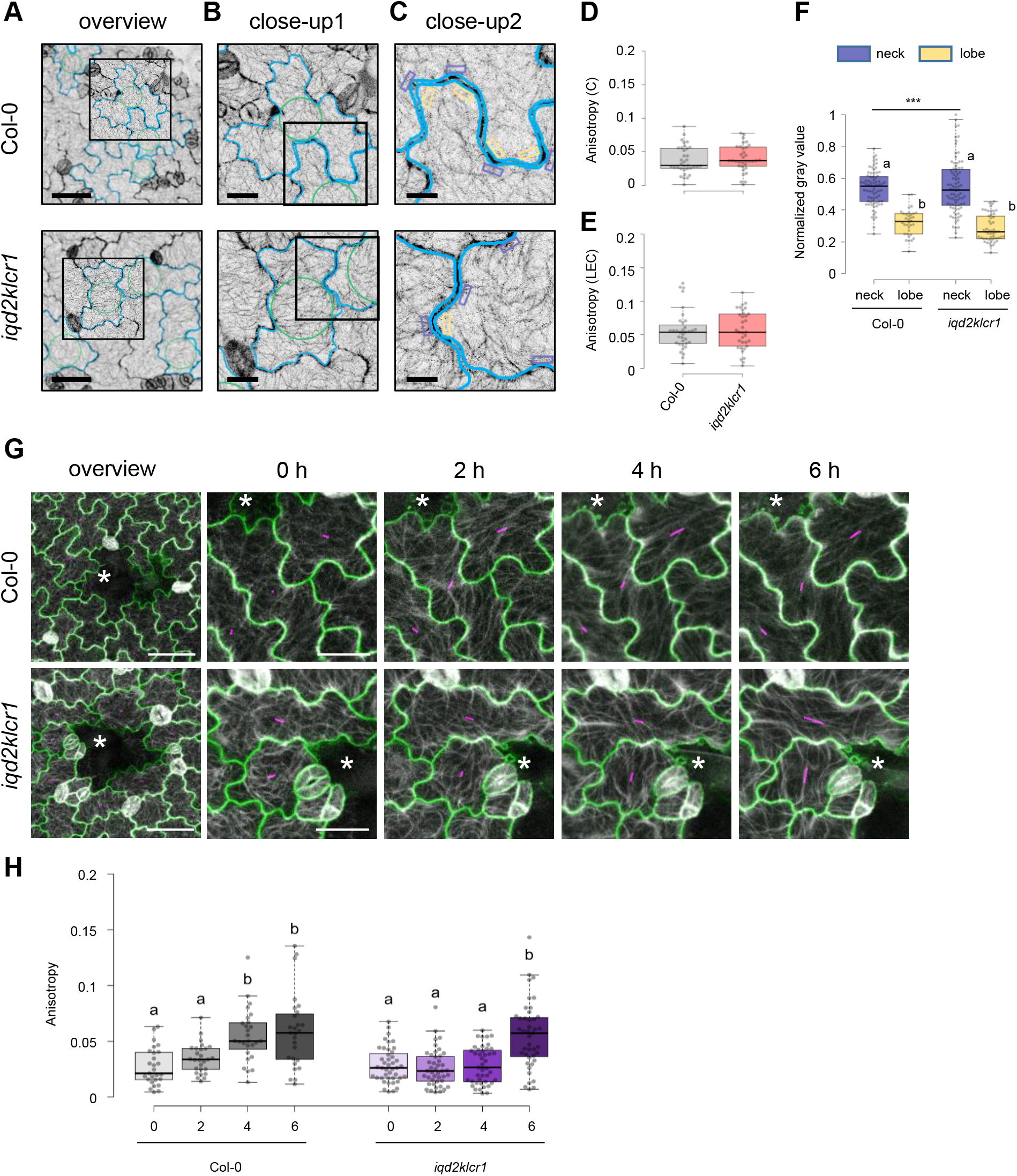
Microtubule organization in pavement cells of *iqd2klcr1* mutants. **A-F**, Microtubule organization in the adaxial side of cotyledon PCs of 5-day-old seedlings from wild type (Col-0) and *iqd2klcr1* expressing the mCh-TUA5 microtubule marker. Representative images of PCs in wild type and *iqd2klcr1* (A), magnification of individual PCs (B), marked by the boxed area in A, and magnification of lobe and neck regions (C), marked by the box in B. Cell contours and the LEC a highlighted in blue and green, respectively. Neck and lobe regions selected for quantitative analysis are marked by purple and yellow boxes, respectively. Quantitative analysis of microtubule anisotropy within complete cells (D) and within LECs (E) in n=35cells (wild type) and n=34 cells (*iqd2klcr1*). Quantification of microtubules in necks and lobes by means of normalized gray values (F) in n=A (neck), 74; B(lobe), 34 (wild type) and n=A(neck), 84; B(lobe), 50 (i*qd2klcr1*). Different letters denote significant statistical differences by ANOVA with TukeyHSD. Statistically significant differences in sample variance are indicated by vertical lines. **G**,**H**, Microtubule reorganization in response to cell ablation in wild type (Col-0) and *iqd2klcr1* immediately after (0 h) laser ablation, and after 2, 4, and 6 hours. Images show cell contours (green) and microtubules (white) in lines expressing the LTi6B-GFP PM marker and the mCh-TUA5 microtubule marker (G). Asterisks mark the ablation side. Pink lines correspond to values of microtubule anisotropy, longer lines indicate higher anisotropy. Quantitiative analysis of microtubule anisotropy with FibrilTool (H) in n= 27 cells from 5 independent seedlings (wild type) and n= 43 cells (*iqd2klcr1*) from 7 independent seedlings. Scale bars, 50 µm (A, G, overview), 25 µm (G), 20 µm (B) and 10 µm (C).

## Acknowledgements

This work was supported by the Deutsche Forschungsgemeinschaft (grants BU2955/2-1 and BU2955/1-2 to K.B.), by the German-Israeli Foundation for Scientific Research and Development (grant G-1482-423.13/2018 to K.B.) and by core funding of the Leibniz Association and Philipps-University Marburg (to K.B.). A.S. acknowledges financial support by the DFG (RTG 2467, project number 391498659 “Intrinsically Disordered Proteins-Molecular Principles, Cellular Functions, and Diseases”, CRC 1423, project number 421152132), INST 271/404-1 FUGG, INST 271/405-1 FUGG), the Federal Ministry for Economic Affairs and Energy (BMWi, ZIM project KK5096401SK0), the region of Saxony-Anhalt, and the Martin Luther University Halle-Wittenberg (Center for Structural Mass Spectrometry). The authors thank Dirk Tänzler for his assistance with MS sample preparation. S.P. was funded by a Villum, three Novo Nordisk, and Danish National Research Foundation grants (25915, 19OC0056076, 20OC0060564, 23OC0086341, DNRF155, respectively). We thank Romina Paradowski, Dipannita Mitra, and Paul Pflug for support with mutant generation and protein expression. We are grateful to Ram Dixit and Arun Sampathkumar for providing *ProKLCR1:mCherry-KLCR1/klcr1* lines and LTi6B-GFP/mCherry-TUA5 marker lines, respectively, and to Petra Dietrich for providing pENTR-CaM2 plasmid.

## Author contributions

J.B, S.K., M.K., P.D., J.P., G.S., and K.B. performed most experiments. Spinning-disc and TIRF microscopy was conducted by L.C., F.R., S.K. and S.P., XL-MS experiments were conducted by C.I. and A.S., and docking and modeling analysis was conducted by C.T. and P.L.K. All authors contributed to data analysis. K.B. conceived the project. J.B., M.K., and K.B. wrote the manuscript draft and all authors read and revised the manuscript.

## Supporting Information

**Supplementary Material and Methods**

**Supplementary Figures S1-S4**

**Supplementary Figure S1** (related to Figure 1): Subcellular localization and dynamics of IQD2-GFP and KLCR1-GFP localization in ProIQD2:IQD2-GFP and ProKLCR1:KLCR1-GFP complementation lines

**Supplementary Figure S2** (related to Figure 2): Purification of KLCR1 and IQD2

**Supplementary Figure S3** (related to Figure 3): Structural predictions and structural determinants of IQD2-KLCR1 binding

**Supplementary Figure S4** (related to Figure 4): Pavement cell shape in wild type, iqd2, klcr1 and iqd2klcr1 mutants

**Supplementary Videos V1-V2**

**Supplementary Video V1:** FRAP analysis of IQD2-GFP

**Supplementary Video V2:** FRAP analysis of KLCR1-GFP

## References

1. Hamant, O. and Haswell, E.S. (2017). Life behind the wall: sensing mechanical cues in plants. BMC Biol 15, 59.

2. Codjoe, J.M., Miller, K. and Haswell, E.S. (2022). Plant cell mechanobiology: Greater than the sum of its parts. Plant Cell 34, 129–145.

3. Monshausen, G.B. and Haswell, E.S. (2013). A force of nature: molecular mechanisms of mechanoperception in plants. J Exp Bot 64, 4663–4680.

4. Baluska, F., Samaj, J., Wojtaszek, P., Volkmann, D. and Menzel, D. (2003). Cytoskeleton-plasma membrane-cell wall continuum in plants. Emerging links revisited. Plant Physiol 133, 482–491.

5. Liu, Z., Persson, S. and Sánchez-Rodríguez, C. (2015). At the border: the plasma membrane-cell wall continuum. J Exp Bot 66, 1553–1563.

6. Schneider, R., Ehrhardt, D.W., Meyerowitz, E.M. and Sampathkumar, A. (2022). Tethering of cellulose synthase to microtubules dampens mechano-induced cytoskeletal organization in Arabidopsis pavement cells. Nat Plants 8, 1064–1073.

7. Hamant, O., Inoue, D., Bouchez, D., Dumais, J. and Mjolsness, E. (2019). Are microtubules tension sensors? Nat Commun 10, 2360.

8. Yan, Y., Sun, Z., Yan, P., Wang, T. and Zhang, Y. (2023). Mechanical regulation of cortical microtubules in plant cells. New Phytol 239, 1609–1621.

9. van Spoordonk, R., Schneider, R. and Sampathkumar, A. (2023). Mechano-chemical regulation of complex cell shape formation: Epidermal pavement cells - A case study. Quant Plant Biol 4, e5.

10. Jacques, E., Verbelen, J.-P. and Vissenberg, K. (2014). Review on shape formation in epidermal pavement cells of the Arabidopsis leaf. Funct Plant Biol 41, 914–921.

11. Gutierrez, R., Lindeboom, J.J., Paredez, A.R., Emons, A.M.C. and Ehrhardt, D.W. (2009). Arabidopsis cortical microtubules position cellulose synthase delivery to the plasma membrane and interact with cellulose synthase trafficking compartments. Nat Cell Biol 11, 797–806.

12. Sampathkumar, A., Krupinski, P., Wightman, R., Milani, P., Berquand, A., Boudaoud, A., Hamant, O., Jönsson, H. and Meyerowitz, E.M. (2014). Subcellular and supracellular mechanical stress prescribes cytoskeleton behavior in Arabidopsis cotyledon pavement cells. eLife 3, e01967.

13. Green, P.B. and King, A. (1966). A mechanism for the origin of specifically oriented textures in development with special reference to Nitella wall texture. Aust J Biol Sci 19, 421–438.

14. Zang, J., Klemm, S., Pain, C., Duckney, P., Bao, Z., Stamm, G., Kriechbaumer, V., Bürstenbinder, K., Hussey, P.J. and Wang, P. (2021). A novel plant actin-microtubule bridging complex regulates cytoskeletal and ER structure at ER-PM contact sites. Curr Biol 31, 1251–1260

15. Liu, Z., Schneider, R., Kesten, C., Zhang, Y., Somssich, M., Zhang, Y., Fernie, A.R. and Persson, S. (2016). Cellulose-Microtubule Uncoupling proteins prevent lateral displacement of microtubules during cellulose synthesis in Arabidopsis. Dev Cell 38, 305– 315.

16. (2011). Evidence for network evolution in an Arabidopsis interactome map. Science 333, 601–607.

17. Ganguly, A., Zhu, C., Chen, W. and Dixit, R. (2020). FRA1 kinesin modulates the lateral stability of cortical microtubules through Cellulose Synthase-Microtubule Uncoupling proteins. Plant Cell 32, 2508–2524.

18. Möller, B., Poeschl, Y., Plötner, R. and Bürstenbinder, K. (2017). PaCeQuant: A tool for high-throughput quantification of pavement cell shape characteristics. Plant Physiol 175, 998–1017.

19. Poeschl, Y., Möller, B., Müller, L. and Bürstenbinder, K. (2020). User-friendly assessment of pavement cell shape features with PaCeQuant: Novel functions and tools. Method Cell Biol 160, 349–363.

20. Yang, B., Stamm, G., Bürstenbinder, K. and Voiniciuc, C. (2022). Microtubule-associated IQD9 orchestrates cellulose patterning in seed mucilage. New Phytol 235, 1096–1110.

21. Ruhnow, F., Persson, S. and Schneider, R. (2023). Noninvasive long-term imaging of the cytoskeleton in Arabidopsis seedlings. Method Mol Biol 2604, 297–309.

22. Bürstenbinder, K., Savchenko, T., Müller, J., Adamson, A.W., Stamm, G., Kwong, R., Zipp, B.J., Dinesh, D.C. and Abel, S. (2013). Arabidopsis Calmodulin-binding protein IQ67-Domain 1 localizes to microtubules and interacts with Kinesin Light Chain-related protein-1. J Biol Chem 288, 1871–1882.

23. Goode, B.L. and Feinstein, S.C. (1994). Identification of a novel microtubule binding and assembly domain in the developmentally regulated inter-repeat region of tau. J Cell Biol 124, 769–782.

24. Cyr, R.J. (1991). Calcium/calmodulin affects microtubule stability in lysed protoplasts. J Cell Sci 100, 311–317.

25. Kölling, M., Kumari, P. and Bürstenbinder, K. (2019). Calcium- and calmodulin-regulated microtubule-associated proteins as signal-integration hubs at the plasma membrane-cytoskeleton nexus. J Exp Bot 70, 387–396.

26. He, L., Hou, Z. and Qi, R.Z. (2008). Calmodulin binding and Cdk5 phosphorylation of p35 regulate its effect on microtubules. J Biol Chem 283, 13252–13260.

27. Zhu, Z.C., Gupta, K.K., Slabbekoorn, A.R., Paulson, B.A., Folker, E.S. and Goodson, H.V. (2009). Interactions between EB1 and microtubules: dramatic effect of affinity tags and evidence for cooperative behavior. J Biol Chem 284, 32651–32661.

28. Mitra, D., Klemm, S., Kumari, P., Quegwer, J., Möller, B., Poeschl, Y., Pflug, P., Stamm, G., Abel, S. and Bürstenbinder, K. (2018). Microtubule-associated protein IQ67 DOMAIN5 regulates morphogenesis of leaf pavement cells in Arabidopsis thaliana. J Exp Bot 70, 529– 543.

29. Feng, X., Pan, S., Tu, H., Huang, J., Xiao, C., Shen, X., You, L., Zhao, X., Chen, Y., Xu, D., et al. (2023). IQ67 DOMAIN protein 21 is critical for indentation formation in pavement cell morphogenesis. J Integr Plant Biol 65, 721–738.

30. Bürstenbinder, K., Möller, B., Plötner, R., Stamm, G., Hause, G., Mitra, D. and Abel, S. (2017). The IQD family of Calmodulin-binding proteins links calcium signaling to microtubules, membrane subdomains, and the nucleus. Plant Physiol 173, 1692–1708.

31. Liang, H., Zhang, Y., Martinez, P., Rasmussen, C.G., Xu, T. and Yang, Z. (2018). The Microtubule-associated protein IQ67 DOMAIN5 modulates microtubule dynamics and pavement cell shape. Plant Physiol 177, 1555–1568.

32. Kumari, P., Dahiya, P., Livanos, P., Zergiebel, L., Kölling, M., Poeschl, Y., Stamm, G., Hermann, A., Abel, S., Müller, S., et al. (2021). IQ67 DOMAIN proteins facilitate preprophase band formation and division-plane orientation. Nat Plants 7, 739–747.

33. Abel, S., Bürstenbinder, K. and Müller, J. (2013). The emerging function of IQD proteins as scaffolds in cellular signaling and trafficking. Plant Signal Behav 8, e24369.

34. Dahiya, P. and Bürstenbinder, K. (2023). The making of a ring: Assembly and regulation of microtubule-associated proteins during preprophase band formation and division plane set-up. Curr Opin Plant Biol 73, 102366.

35. Abel, S., Savchenko, T. and Levy, M. (2005). Genome-wide comparative analysis of the IQD gene families in Arabidopsis thaliana and Oryza sativa. BMC Evol Biol 5, 72.

36. Dahiya, P., Haluška, S., Buhl, J., Kölling, M., Papsdorf, S., Zehnich, D., Machalett, K., Pfeiffer, P., Stamm, G., Potocký, M., et al. (2023). Origin and evolution of IQD scaffolds and assembled protein complexes in plant cell division. SSRN 10.2139/ssrn.4655235.

37. Sugiyama, Y., Wakazaki, M., Toyooka, K., Fukuda, H. and Oda, Y. (2017). A novel plasma membrane-anchored protein regulates xylem cell-wall deposition through microtubule-dependent lateral inhibition of Rho GTPase domains. Curr Biol 27, 2522–2528.

38. Li, Y., Huang, Y., Wen, Y., Wang, D., Liu, H., Li, Y., Zhao, J., An, L., Yu, F. and Liu, X. (2021). The domain of unknown function 4005 (DUF4005) in an Arabidopsis IQD protein functions in microtubule binding. Journal Biol Chem 297, 100849.

39. Covarrubias, A.A., Cuevas-Velazquez, C.L., Romero-Pérez, P.S., Rendón-Luna, D.F. and Chater, C.C.C. (2017). Structural disorder in plant proteins: where plasticity meets sessility. Cell Mol Life Sci 74, 3119–3147.

40. Covarrubias, A.A., Romero-Pérez, P.S., Cuevas-Velazquez, C.L. and Rendón-Luna, D.F. (2020). The functional diversity of structural disorder in plant proteins. Arch Biochem Biophys 680, 108229.

41. Perez-Riba, A. and Itzhaki, L.S. (2019). The tetratricopeptide-repeat motif is a versatile platform that enables diverse modes of molecular recognition. Curr Opin Struct Biol 54, 43– 49.

42. Allan, R.K. and Ratajczak, T. (2011). Versatile TPR domains accommodate different modes of target protein recognition and function. Cell Stress Chaperones 16, 353–367.

43. Iacobucci, C., Götze, M., Ihling, C.H., Piotrowski, C., Arlt, C., Schäfer, M., Hage, C., Schmidt, R. and Sinz, A. (2018). A cross-linking/mass spectrometry workflow based on MS-cleavable cross-linkers and the MeroX software for studying protein structures and protein-protein interactions. Nat Protoc 13, 2864–2889.

44. Hage, C., Falvo, F., Schäfer, M. and Sinz, A. (2017). Novel concepts of MS-cleavable cross-linkers for improved peptide structure analysis. J Am Soc Mass Spectrom 28, 2022– 2038.

45. Jumper, J., Evans, R., Pritzel, A., Green, T., Figurnov, M., Ronneberger, O., Tunyasuvunakool, K., Bates, R., Žídek, A., Potapenko, A., et al. (2021). Highly accurate protein structure prediction with AlphaFold. Nature 596, 583–589.

46. van Zundert, G.C.P., Rodrigues, J.P.G.L.M., Trellet, M., Schmitz, C., Kastritis, P.L., Karaca, E., Melquiond, A.S.J., van Dijk, M., Vries, S.J. de and Bonvin, A.M.J.J. (2016). The HADDOCK2.2 web server: User-friendly integrative modeling of biomolecular complexes. J Mol Biol 428, 720–725.

47. Lu, Q., Li, J., Ye, F. and Zhang, M. (2015). Structure of myosin-1c tail bound to calmodulin provides insights into calcium-mediated conformational coupling. Nat Struct Mol Biol 22, 81–88.

48. Eng, R.C., Schneider, R., Matz, T.W., Carter, R., Ehrhardt, D.W., Jönsson, H., Nikoloski, Z. and Sampathkumar, A. (2021). KATANIN and CLASP function at different spatial scales to mediate microtubule response to mechanical stress in Arabidopsis cotyledons. Curr Biol 31, 3262–3274.e6.

49. Kumar, M., Carr, P. and Turner, S.R. (2022). An atlas of Arabidopsis protein S-acylation reveals its widespread role in plant cell organization and function. Nat Plants 8, 670–681.

50. Haley, A., Russell, A.J., Wood, N., Allan, A.C., Knight, M., Campbell, A.K. and Trewavas, A.J. (1995). Effects of mechanical signaling on plant cell cytosolic calcium. Proc Natl Acad Sci U S A 92, 4124–4128.

51. Trewavas, A. and Knight, M. (1994). Mechanical signalling, calcium and plant form. Plant Mol Biol 26, 1329–1341.

52. Burgoyne, T., Patel, S. and Eden, E.R. (2015). Calcium signaling at ER membrane contact sites. Biochim Biophys Acta 1853, 2012–2017.

53. Hepler, P.K. (2016). The cytoskeleton and its regulation by calcium and protons. Plant Physiol 170, 3–22.

