## Supplementary Data for "IQD2 recruits KLCR1 to the membrane-microtubule nexus to promote cytoskeletal mechano-responsiveness in leaf epidermis pavement cells"

| REAGENT or RESOURCE | SOURCE | IDENTIFIER |
|---|---|---|
| Antibodies | | |
| monoclonal mouse HRP-conjugated α-His antibody | Miltenyi Biotec | 130-092-785 |
| monoclonal mouse α-GST antibody | St. John's Laboratory | STJ96909 |
| monoclonal mouse α-S-tag antibody | Sigma Aldrich | SAB2702204 |
| polyclonal HRP-conjugated α-mouse IgG | Sigma Aldrich | A9044 |
| Bacterial and virus strains | | |
| *Escherichia coli* TOP10 | N/A | |
| *Escherichia coli* KRX | N/A | |
| *Agrobacterium tumefaciens* GV3101 pKan | N/A | |
| Chemicals, peptides, recombinant proteins | | |
| MES | Roth | Cat#4256.2 |
| SD | Sigma | Cat#436143 |
| Plant-agargel | Sigma | Cat#P8169 |
| Kanamycin | Roth | Cat#T832.2 |
| Rifampicin | Roth | Cat#4163.1 |
| Gentamycin | Roth | Cat#0233.4 |
| Spectinomycin | Sigma | Cat#S4014 |
| PIPES | Roth | Cat#9156.2 |
| EGTA | Roth | Cat#3054.3 |
| Triton X-100 | Roth | Cat#3051.3 |
| BSA | Roth | Cat#8076.1 |
| propidium iodide | Sigma | Cat#P4170 |
| EcoRI | Thermo Fischer Scientific | Cat#ER0271 |
| XbaI | Thermo Fischer Scientific | Cat#ER0681 |
| XhoI | Thermo Fischer Scientific | Cat#ER0695 |
| Critical commercial assays | | |
| Gateway BP Clonase II | Thermo Fischer Scientific | Cat#11789100 |
| Gateway LR Clonase II | Thermo Fischer Scientific | Cat#11791100 |
| Roche cOmplete Protease Inhibitor Cocktail | Merck | Cat#04693116001 |
| Microtubule Spin Down Kit | Cytoskeleton Ink. | Cat#BK029 |
| Phusion Polymerase | Thermo Fischer Scientific | Cat#F530S |
| Experimental models: cell lines | | |
| PJ694a yeast strain | | |
| Experimental models: organisms/strains | | |
| *Arabidopsis:* Col-0 | NASC | |
| Arabidopsis: *klcr1* | Zang et al. | SAIL_335_B08 |
| Arabidopsis: *klcr2* | Zang et al. | SALK_148296 |
| Arabidopsis: *iqd2* | Ganguly et al. | GK_608F07-021865 |
| Arabidopsis: *klcr1klcr2* | This paper | |
| Arabidopsis: *iqd2klcr1* | This paper | |
| Arabidopsis: *ProIQD2:IQD2-GFP* in *iqd2* | Zang et al. | |
| Arabidopsis: *ProIQD2:IQD2-GFP* in *iqd2klcr1* | This paper | |
| Arabidopsis: *ProKLCR1:KLCR1-GFP* in *klcr1* | Zang et al. | |
| Arabidopsis: *ProKLCR1:KLCR1-GFP* in *iqd2klcr1* | This paper | |
| Arabidopsis: *ProKLCR1:mCherry-KLCR1* in *klcr1* | Ganguly et al. | |
| Arabidopsis: *Pro35S:mCherry-TUA5* | Gutierrez et al. | |
| Arabidopsis: *Pro35S:mCherry-TUA5/ProIQD2:IQD2-GFP* in *iqd2* | This paper | |
| Arabidopsis: *Pro35S:mCherry-TUA5/ProKLCR1:KLCR1-GFP* in *klcr1* | This paper | |
| Arabidopsis: *Pro35S:mCherry-TUA5/ProKLCR1:KLCR1-GFP* in *iqd2klcr1* | This paper | |
| Arabidopsis: *ProIQD2:IQD2-GFP/ProKLCR1:mCherry-KLCR1* in *iqd2klcr1* | This paper | |
| Arabidopsis: *Pro35S:mCherry-TUA5* in *iqd2klcr1* | This paper | |
| Arabidopsis: *Pro35S:mCherry-TUA5/Pro35S:LTi6B-GFP* | Eng et al. | |
| Arabidopsis: *Pro35S:mCherry-TUA5/Pro35S:LTi6B-GFP* in *iqd2klcr1* | This paper | |
| Oligonucleotides | | |
| Primers for amplifying coding sequences, see Table S1 and Table S2 | | |
| Primers for genotyping, see Table S1 and Table S2 | | |
| Recombinant DNA | | |
| Entry gateway vecor: pENTR3C | | |
| Entry gateway vecor: pDONR221 | | |
| Entry gateway vector: pDONR207 | | |
| Empty gateway destination vector: pB7FWGF2 | Karimi et al., 2002 | |
| Empty gateway destination vector: pGWB455 | Addgene | |
| Empty gateway destination vector: pDEST32 | | |
| Empty gateway destination vector: pDEST22 | Addgene | |
| Empty gateway destination vector: pDEST15 | Addgene | |
| Empty gateway destination vector: pDEST SUMO | Addgene | |
| pET41a(+) | | |
| Software and algorithms | | |
| ImageJ | https://imagej.net/software/fiji/ | |
| Inkscape | https://inkscape.org/ | |

| | |
|---|---|
| Pymol | https://www.pymol.org |
| PaCeQuant | https://www.quantitative-plant.org/software/pacequant |
| MiToBo | MiToBo (uni-halle.de) |
| FibrilTool | https://www.quantitative-plant.org/software/fibriltool |
| HADDOCK | https://rascar.science.uu.nl/ |
| Alphafold | https://alphafold.ebi.ac.uk |

#### RESOURCE AVAILABILITY

##### Lead contact

##### Materials availability

Unique materials used in this study will be freely available.

#### EXPERIMENTAL MODEL AND SUBJECT DETAILS

The wild type Columbia-0 (Col-0) was used in all experiments. The T-DNA insertion mutants and transgenic lines in this background are detailed in the Key resources table.

#### METHOD DETAILS

##### Plant material and transformations

*Arabidopsis* wild-type seeds (Col-0) and T-DNA insertion mutants were obtained through the Nottingham Arabidopsis stock center. The *klcr1* (SAIL_335_B08) and *iqd2* (GK_608F07-021865) T-DNA insertion mutants and transgenic complementation lines in these backgrounds were established in a previous study [1]. Establishment of the *klcr2* (SALK_148296C) T-DNA insertion line, also referred to as *cmu2*, and of *ProIQD2:IQD2-GFP/iqd2*, *ProKLCR1:KLCR1-GFP/klcr1*, and *ProKLCR1:mCh-KLCR1/klcr1* lines is described previously [1,2]. As markers, the mCh-TUA5 line [3] and the LTi6B-GFP/mCh-TUA5 dual marker line [4] were used. All markers were introduced into mutant lines by crossing to avoid positional effects on marker gene expression. Homozygosity of mutants was validated by PCR-based genotyping. Primers used for genotyping and primer combinations are listed in Table S1 and Table S2, respectively. Presence of marker constructs was evaluated by microscopic analysis. Seeds were surface-sterilized with chlorine gas, and after 2 d of stratification at 4°C grown vertically on *Arabidopsis thaliana Salts* (ATS) media, 0.5% (w/v) agar gel and 1% (w/v) sucrose in a growth chamber with a 16h (22°C):8h(18°C) light:dark regime [1]. *N. benthamiana* plants were grown at 21 °C on soil under long-day (16h light, 8h dark) conditions. For transient expression assays in *N. benthamiana*, leaves of four-week-old plants were co-infiltrated with *Agrobacterium tumefaciens* GV3101 pMP90RK harboring plasmids and the silencing suppressor p19 in a 1:1 ratio. Bacterial cultures were adjusted to an optical density at 600 nm of 0.5 using infiltration buffer, and *N. benthamiana* leaves were pressure infiltrated through the abaxial epidermis. Transiently transformed *N. benthamiana* leaves were imaged two days after infiltration using confocal laser scanning microscopy.

##### Plasmid generation

DNA sequence information of IQD2 (AT5G03040.1), KLCR1 (AT4G10840.1), and CaM2 (AT2G41110.1) was obtained from The Arabidopsis Information Resource. Gateway-compatible ENTR Clones harboring full length coding sequences of IQD2, KLCR1, and CaM2 were established in previous studies [5–7]. IQD2 variants were amplified from IQD2-pENTR and cloned into pDONR207 or pDONR221 using Gateway BP clonase II (Invitrogen), or into pENTR3C using restriction cloning using the primers and primer combinations listed in Table S1 and Table S2. To generate GFP- and RFP-fused variants, IQD2 and KLCR1 were mobilized into pB7WGF2 [8] and pGWB455 [9], respectively, using LR clonase II (Invitrogen). For yeast two-hybrid assays, GAL4-DNA binding domain (DBD) and activation domain (AD) fusions were generated by mobilization into pDEST32 and pDEST22, respectively [10]. For recombinant expression, IQD2 and IQD2 variants were mobilized into pDEST-SUMO [11]. KLCR1 and CaM2 were cloned into vector pET41a(+) (Novagen) using circular polymerase extension cloning (CPEC) according to [12]. GST-KLCR1, used in *in vitro* pull-down experiments with HS-IQD2 variants, was generated by mobilization into pDEST15 (Invitrogen).

#### Fluorescence microscopy and quantitative image analysis

Fluorescence images (Fig.2E; Fig.3C,D; Fig.4A-C,G; Fig.S1A; Fig.S4A) were recorded using an LSM780, LSM880, LSM980 or LSM900 inverted confocal laser scanning microscope, using a 40x water immersion objective or a 10x objective as described previously [13]. GFP fluorescence was excited with a 488 nm laser, and detected with a band-pass filter between 493 and 535 nm. RFP/mCherry/propidium iodide (PI) fluorescence were excited with a 561 nm laser and detected between 570 and 633 nm. For colocalization analyses, channels were recorded in the sequential scanning mode. Z-stacks were generated with the corresponding optimal interval suggested by Zeiss' Zen software. For spinning disc microscopy (Fig.1A,B,E,F,I-L; Fig.2A-D; Fig.S1G-J), seedlings were grown in small imaging chambers as described in [14] Fluorescence images were recorded using a custom-built microscope system (Intelligent Imaging Innovations), consisting of a Nikon Eclipse Ti-E2 equipped with a Yokogawa CSU-W1 spinning disc unit (50µm pinhole disk; Yokogawa) and 2 Andor iXon Life 888 EMCCD cameras (Oxford Instruments), with a 100x oil immersion objective (Nikon CFI60 Apo 100x/1.49NA). GFP fluorescence was excited with a 488 nm laser using a 405/488/561/640 quad-band dichroic, and detected with a 525/30 band-pass filter. mCherry fluorescence was excited with a 561 nm laser using a 405/488/561/640 quad-band dichroic, and detected with a 617/73 band-pass filter. For subcellular localization, either 100 time-lapse images of a single z-plane were acquired in streaming mode using Slidebook (Intelligent Imaging Innovations) using a 200ms exposure time and single images were generated in FIJI by averaging the intensities of all frames using the "image/stacks/Z project/" option (Fig.1A,B,E,F) or several z-planes were acquired with 10 frame averaging (100ms exposure time each) and single images were generated in FIJI using the maximum projection option (Fig.2A-F).. For dynamics, fluorescence images of a single z-plane were acquired by averaging 10 frames with 100ms exposure time each at 4s intervals (Fig.1I-L). Fluorescence Recovery after Photobleach (FRAP) experiments were conducted in PCs of the adaxial side of cotyledons from eight-day.old seedlings. Bleaching was performed on a rectangular ROI of 13x13 µm after the 3rd time-point with 100% laser intensity. Fluorescence recovery was imaged at a single z-plane by averaging 10 frames with 100ms exposure time each at 4s intervals (Fig.S1G,H). The same seedlings were imaged on a TIRF microscope (Fig.1M) by transferring the imaging chambers between two systems. Fluorescence TIRF images were recorded using a Zeiss Elyra PS.1 microscope (Zeiss) with a 100x oil immersion objective (Zeiss alpha Plan-Apochromat 100x/1,46 Oil), 1.6x additional magnification (Zeiss Optovar) and Andor iXon+ 897 EMCCD cameras (Oxford Instruments). GFP fluorescence was excited with a 488 nm laser using a 488 dichroic, and detected with a 535/80 band-pass filter. mCherry fluorescence was excited with a 561 nm laser using a 561 dichroic, and detected with a 610/80 band-pass filter. For colocalization, 50 time-lapse images of a single z-plane were acquired in streaming mode using ZEN (Zeiss) using a 200ms exposure time. Single images were generated in ZEN by averaging the intensities of all frames. For colocalization studies, in tobacco transient expression system, around 100 cells were evaluated that expressed both the proteins and the percentage of cells for the specific localization was calculated.

Images were analyzed and processed using Fiji/ImageJ [15]. Maximum intensity projections of z-stacks were generated using the "image/stacks/Z project/" option. Fluorescence profiles were generated along a line inserted into the image, fluorescence intensities read out via the "analyze/plot profile" option, and analyzed via MS Excel. Fluorescence intensities were normalized to the highest intensity determined per plot profile. The PCC was calculated in Excel using the gray values of GFP signal as matrix 1 and the gray values of RFP signal as matrix 2. Time series images were loaded into ImageJ. A line was drawn along a selected microtubule. Kymograph was created via Reslice tool of Fiji.

#### Quantification of PC shape, microtubule organization, and ablation experiments

PC shape was analyzed in the adaxial side of cotyledons of five-day-old seedlings using the PaCeQuant tool as described previously [16–19]. Cell contours were visualized by PI staining, and dissected cotyledons were placed on microscopy slides in liquid ATS media. Single optical sections of the anticlinal walls were acquired by confocal laser scanning microscopy with a 20x objective. Automatic cell segmentation was conducted with the “segmentation” operation in PaCeQuant, segmentation results were manually corrected using the LabelImageEditor tool, and shape features were calculated using PaCeQuant’s “feature extraction” option. Statistical analysis and graphical visualization in violin plots were performed using the R script “PaCeQuantAna”. Heatmaps of feature distributions were generated with MiToBo’s supplementary FeatureColorMapper [19,20].

Quantification of microtubule organization and ablation experiments were performed with wild type and *iqd2klcr1* seedlings expressing LTi6B-GFP/mCH-TUA5 dual marker [4]. Microtubule anisotropy was quantified with FibrilTool [21] in cotyledons of five-day-old seedlings. As input, maximum projections of z-stacks covering the upper half of pavement cells from the outer periclinal to the anticlinal wall were used and anisotropy was calculated within ROIs of whole cells or the corresponding LECs, extracted with PaCeQuant. Microtubule enrichment in the lobe and neck regions was performed by selecting rectangular ROIs of the same size at necks and lobes as described in as described in [17]. The gray value for each lobe and neck region was measured in ImageJ.

Ablation experiments were performed with four-day-old seedlings. Seedlings were transferred on microscopy slides covered with 50 µl low melting agar. Laser ablations were performed with a Zeiss LSM900 confocal microscope using the ZEN software with a circular ROI (diameter 31 µm) as described in [22]. Z-stacks of pavement cells surrounding the ablation site were recorded immediately after ablation (0h), and in 2 h intervals after 2, 4, and 6 h and microtubule anisotropy was measured with Fibril tool within the upper periclinal to anticlinal cell surface.

#### Yeast two-hybrid assays

For yeast two-hybrid assays, the coding sequences of IQD2 and IQD2 variants were mobilized into the bait pDEST32 vector to generate protein fusions with the GAL4 DNA binding domain (DBD). CDS of KLCR1 and CaM2 were mobilized into the prey vector pDEST22 to generate fusion proteins with the GAL4 activation domain (AD). Bait and prey vectors to be tested for interaction were co-transformed into *Saccharomyces cerevisiae* strain PJ69-4a using the standard LiOAc method [23]. Briefly, 10 µl of the yeast were spotted on plates with vector-selective media (double drop out (DDO) media: SD/-Leucin/-Tryptophan) and were grown for 3 days at 29°C. Two to three independent colonies of each transformation were inoculated in 1x TE buffer and incubated at 29°C overnight. The next day, OD600 was adjusted to 0.5 -1 and 10 µl each of the colonies were spotted on DDO and interaction-selective media (triple drop out (TDO) media: SD/-Leucin/-Tryptophan/-Histidin). Pictures were taken three days after spotting. SD plates were prepared with SD media and 1.6% (w/v) Bactoagar.

#### Recombinant protein expression and purification

All proteins and protein fragments used in this study were expressed in *E. coli* KRX cells. HS-IQD2 was co-expressed with GH-CaM2 to improve solubility [24]. Protein expression was induced with 1.4 mM rhamnose and 0.5 mM IPTG and proteins were expressed at 16 °C for

20 to 24 hours. For expression of GST, protein expression was conducted for 3 to 4 hours at 37 °C. GH-S-tag-KLCR1 was purified using tandem affinity purification, combining immobilized metal-ion affinity chromatography (IMAC) via Ni-NTA with affinity purification via GSH-agarose. Affinity purification was followed by on-column tag-removal by thrombin-cleavage and final purification and thrombin removal by gel filtration on a Superdex 200 column using an ÄKTA FPLC system. Proteins were eluted from Ni-NTA with 200 mM imidazole and from GSH-agarose with 10 mM GSH. GSH and imidazole were removed by overnight dialysis into elution buffer without GSH or imidazole. Cell lysis and IMAC were carried out in Tris-lysis, -wash, and -elution buffer (Tris-base buffer: 50 mM Tris-HCl pH 8.0, 200 mM NaCl; Tris-lysis buffer: Tris-base buffer, 5 mM imidazole, 1 mM PMSF, 1x Roche cOmplete protease inhibitor cocktail, 0.1 % (v/v) Tween20; Tris-wash buffer: Tris-base buffer, 10, 25, or 50 mM imidazole, 1 mM $MgCl_2$ and 5 mM ATP in wash 3; Tris-elution buffer: Tris-base buffer, 250 mM imidazole, 10 % (v/v) glycerol). For pET41a(+)-KLCR1, pEXP-SUMO-IQD2$_{C\text{-term}}$, and pEXP-SUMO-IQD2$_{C\text{-half}}$ IMAC included a high-salt wash with 1 M NaCl during the second washing step. For GSH-affinity purification, the Tris-buffer was supplemented with 1 mM DTT or DTE. The Tris-buffer was exchanged to HEPES- or PIPES-buffer (HEPES buffer: 20 mM HEPES-KOH pH 7.5, 50 mM NaCl, 10 % (v/v) glycerol, 0.2 mM TCEP; PIPES buffer: 80 mM PIPES-KOH pH 7.5, 50 mM NaCl, 2 mM $MgCl_2$, 10 % (v/v) glycerol, 0.2 mM TCEP) during the second washing step of the GSH-affinity purification. For co-purification of HS-IQD2 and GH-CaM2, purification buffers were supplemented with 1 mM $CaCl_2$. The $CaCl_2$-concentration was gradually decreased to 100 µM during GSH-affinity purification. Purified proteins were quantified using BSA calibration curves (IQD2/CaM2) and UV/vis quantification (KLCR1).

#### *In vitro* pulldown assays and microtubule spin-down assays

Microtubule spin-down assays were carried out using the “Microtubule Binding Protein Spin-Down Assay Biochem Kit” (mostly according to manufacturer instructions. Centrifugation was conducted at 20,000 g, the amount of tubulin per reaction was halved, and spin-downs were carried out in the presence of 100 µM $CaCl_2$. Proteins in supernatant and pellet fractions were detected by trichloroethanol (TCE) in-gel visualization or Coomassie Brilliant Blue (CBB) staining. To discriminate between HS-IQD2 and S-tag-KLCR1 with similar calculated molecular masses of ~65 and ~70 kDa, respectively, and a similar migration behavior at ~70 kDa in denaturing SDS-PAGE, proteins were detected by Western blot. Quantification of protein levels in the pellet and supernatant was performed using ImageJ, based on CBB-stained gels or in-gel visualization with TCE. For S-tag-KLCR1 in the presence of HS-IQD2, quantification was based on antibody detection of both proteins. Proteins were detected by Western blotting, using an HRP-conjugated α-His antibody, and a primary mouse monoclonal α-S-tag antibody, combined with a secondary HRP-conjugated anti-mouse antibody. Images were imported into ImageJ and converted to an 8-bit grayscale format. Individual lanes were selected using the rectangular selection tool, followed by the Analyze > Gels > Select First/Second Lane command. Lane profiles were then generated using the Analyze > Gels > Plot Lanes function. The area under each peak was measured, and for each protein, the areas of the pellet and supernatant fractions were summed to represent the total protein amount and to calculate the percentage of protein in the pellet and supernatant. In the case of S-tag-KLCR1 in the presence of HS-IQD2, the relative amount of S-tag-KLCR1 in the pellet was normalized to the largest peak area observed for S-tag-KLCR1 across the two spin-down assays and both fractions. Pull-down assays were carried out with HS-IQD2 fragments and GST-KLCR1. HS-IQD2 fragments were immobilized on Ni-NTA agarose and incubated with GST-KLCR1 or GST alone as control over night at 4 °C. Bead-bound proteins were boiled in Laemmli buffer at 95 °C for 5 minutes. Proteins were detected by Western blotting, using an HRP-conjugated α-His antibody, and a primary mouse α-GST combined with a secondary HRP-conjugated α-mouse IgG. For detection the ECL detection Kit from Cytiva (pico and femto in a 1:1 ratio) was used (Cytiva AmershamTM ECL SelectTM Western Blotting Detection Reagent).

#### XL-MS experiments

XL-MS was performed according to [25]. Proteins were chemically cross-linked using a 30x molar excess of the amine-reactive, MS-cleavable cross-linker DSBU in HEPES-buffer supplemented with 1 mM $CaCl_2$. The concentration of KLCR1 was adjusted to 5 µM. Eluates from co-purification of IQD2 and CaM2 were used undiluted after dialysis. Proteins were incubated at RT for 30 minutes prior to the addition of DSBU and cross-linking reactions were stopped with 4x Laemmli buffer after two hours at RT. Cross-linking reaction mixtures were analyzed via SDS-PAGE (10% resolving gel). Bands corresponding to cross-linked complexes were excised and in-gel digestion was performed with AspN and trypsin following our protocol described in [25]. Resulting peptide mixtures were subjected to LC-MS/MS analysis using an UltiMate 3000 RSLC nano-HPLC system (Thermo Fisher Scientific) coupled to a timsTOF Pro mass spectrometer equipped with CaptiveSpray source (Bruker Daltonik) [26]. Peptides were trapped on a C18 pre-column (Acclaim PepMap 100, 300 µm x 5 mm, 5 µm, 100 Å (Thermo Fisher Scientific) and separated on a µPAC 50 C18 column (PharmaFluidics). After trapping, peptides were eluted by a 90 min water-ACN gradient from 3 % (v/v) to 35 % (v/v) ACN at a flow rate of 600 nl/min. For MS data acquisition, parameters were used as described in [26].
Cross-links were identified via MeroX search (version 2.0.1.7) [27] against the sequences of the KLCR1, CaM2, and IQD2 constructs using the following settings: Proteolytic cleavage C-terminal at Lys and Arg, and N-terminal at Asp and Glu; peptide lengths of 4 to 50 amino acids; modifications: alkylation of Cys by IAA, oxidation of Met; cross-linker specificity Lys, Ser, Thr, Tyr, N-terminus; search algorithm RISEUP mode; precursor mass accuracy of 15 ppm; fragment ion mass accuracy of 20 ppm; signal-to-noise ratio ≥ 1.5; precursor mass correction enabled; 10 % intensity as pre-score cutoff; 1 % false-discovery rate cutoff; minimum score of 60. The resulting .csv files were visualized as circle plots using xiView (https://www.xiview.org/xiNET_website/index.php).

#### *In silico* modelling of cross-linked protein-complexes

Monomeric models of KLCR1, IQD2, and CaM2 were generated using a local installation of Alphafold v2.2 with default parameters [https://www.nature.com/articles/s41592-022-01488-1]. The HADDOCK web server 2.2 (guru interface) [https://www.sciencedirect.com/science/article/pii/S0022283615005379] was used for docking, and generating structures of CaM2-IQD2-KLCR1-complexes. Cross-links were defined as unambiguous distance restraints between the Cα-atoms of the cross-linked amino acid residues with a permitted distance of 0 to 30 Å. Regions with low prediction confidence, as indicated by the plDDT score, were removed prior docking. Euklidean distances were determined by extracting the coordinates of each Cα atom from the .pdb files generated by AlphaFold or HADDOCK. A cross-link was considered satisfied with a Cα-distance d below 30 Å. The final model was generated by superposition of the best ranked docking solution of IDQ2-KLCR1 and the AlphaFold2 model of CaM2-IQD2.

#### *In silico* disorder and charge prediction

Disorder predictions for IQD2 and KLCR1 were carried out by submitting full-length amino acid sequences to disorder prediction servers PONDR, IUPRED, PrDOS, MFDp2, and Disopred with default parameters. Disorder consensus was calculated as the average disorder propensity per residue. A disorder propensity > 0.5 was considered indicative of intrinsic disorder. Charge plots were generated by submitting full-length amino acid sequences to the EMBOSS charge tool and plotting the resulting charge predictions per amino acid residue in MS Excel.

### QUANTIFICATION, STATISTICAL ANALYSIS AND DATA VISUALIZATION

For box plots, data are presented as median (center line); upper and lower quartiles (box limits); interquartile range (whiskers); outliers (points); individual data points are shown in bee swarms. For violin plots, means and medians are represented by crosses and circles, respectively. Thick vertical black lines represent standard deviation (SD); thin black lines represent 95% confidence interval; the width of the violin represents local distribution of values along the y axes. Box plots were generated in R with BoxplotR (http://shiny.chemgrid.org/boxplotr/). The violin plots were generated in R with PaCeQuantAna (https://mitobo.informatik.uni-halle.de/index.php/Applications/PaCeQuantAna). Labeling of plots was added in Inkscape. Statistical analysis of data was performed by one-way ANOVA with a post-hoc Tukey's honestly significant difference test (https://astatsa.com/OneWay_Anova_with_TukeyHSD). To test for differences in sample variance, Levene's test was used (https://www.socscistatistics.com/tests/levene/default.aspx). For statistical analysis of pavement cell shape features, the non-parametric Kruskal-Wallis test with a post-hoc Dunn's test was applied using the R package PaCeQuantAna. Sample sizes and specific descriptions of statistical tests are indicated in the figure legends.

**Supplemental Information**

### IQD2 recruits KLCR1 to the membrane-microtubule nexus to promote cytoskeletal mechano-responsiveness in leaf epidermis pavement cells

Jonas Buhl, Malte Kölling, Sandra Klemm, Leia Colin, Felix Ruhnow, Christian Ihling, Christian Tüting, Pradeep Dahiya, Jacqueline Patzsch, Gina Stamm, Andrea Sinz, Panagiotos L. Kastritis, Staffan Persson, Katharina Bürstenbinder

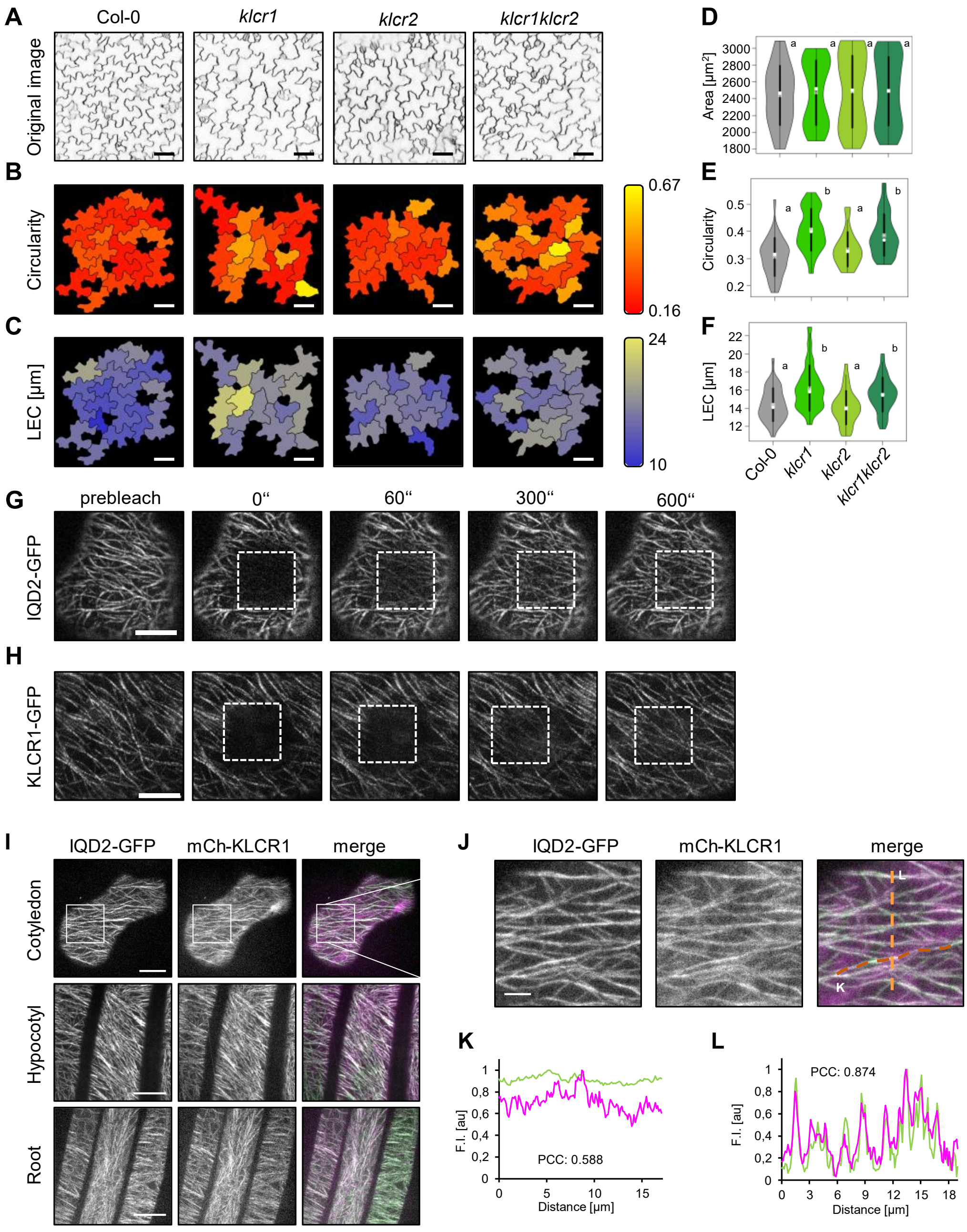


**Figure S1 (related to Figure 1): Subcellular localization and dynamics of IQD2-GFP and KLCR1-GFP localization in *ProIQD2:IQD2-GFP* and *ProKLCR1:KLCR1-GFP* complementation lines**
**A-F,** Pavement cell (PC) shape in the adaxial side of cotyledons of 5-day-old seedling of wild type (Col-0), *klcr1, klcr2* and *klcr1klcr2* mutants. Representative images of PC morphology (A); cell outlines were visualized with PI, images are single optical sections. Heatmaps of circularity (B) and largest empty circle (LEC). Violin plots of feature distributions for area (D), circularity (E), and LEC (F) from n=36 (Col-0), n=24 (*klcr1*), n=33 (*klcr2*) and n=43 (*klcr1klcr2*) cells, analyzed in five cotyledons from five seedlings each. Different letters denote significant statistical differences ($p < 0.001$, Kruskal-Wallis. Post-hoc Dunn's test). **G,H,** FRAP analysis of IQD2-GFP (G) and KLCR1-GFP (H) signals. Images of surface section of cotyledons of 8-day-old seedlings. **I-L,** Subcellular co-localization of IQD2-GFP and mCh-KLCR1 in cotyledons, hypocotyl and root cells of 10-day-old seedlings (I). Magnification of boxed area of cotyledon section (J). Quantitative analysis of co-localization by Pearson correlation coefficients (PCC) of fluorescence intensity profiles from IQD2-GFP (green) and mCh-KLCR1 (magenta) along a selected microtubule (K, horizontal line in J) and across multiple microtubules (L, vertical line in J). Scale bars, 50 µm (A-C); 10 µm (G,H); 15 µm (I); 5 µm (J).

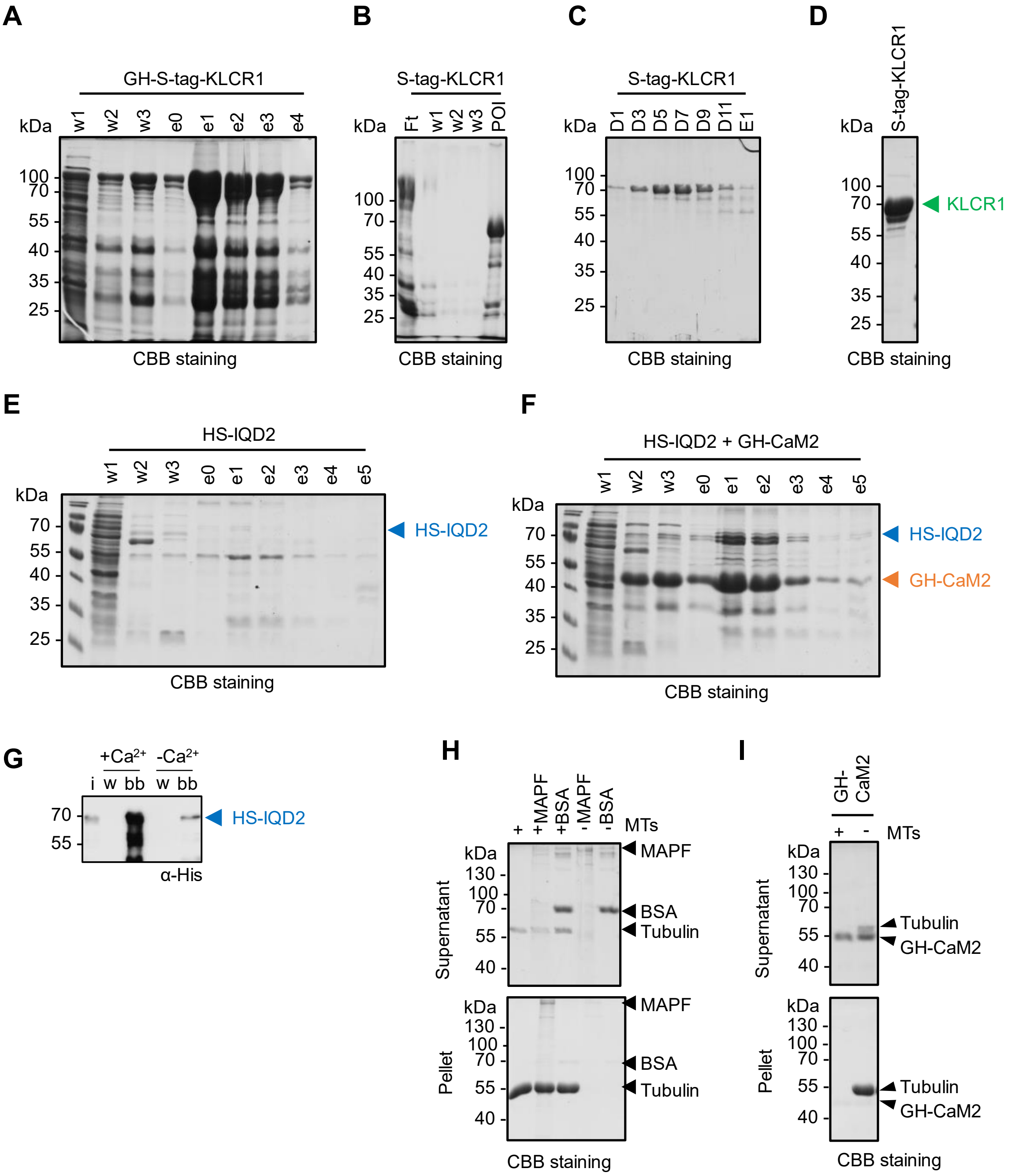


**Figure S2 (related to Figure 2): Purification of KLCR1 and IQD2**

**A-D:** Coomassie-stained SDS-PAGE of purification of GST-His-S-tag-KLCR1 via IMAC (A) followed by GST-His-tag removal via thrombin-cleavage (B), and gel filtration (C), resulting in pure S-tag-KLCR1 protein (D). **E,F,** Coomassie-stained SDS-PAGE of purification of His-SUMO-IQD2 expressed alone (E), and in co-expression and –purification with GST-His-CaM2 (F) via IMAC. w, wash fraction; e, elution fraction. **G,** Western blot of CaM-pulldown of HS-IQD2 in the presence and absence of Calcium. I, input; w, wash; bb, bead-bound fraction **H,I,** Coomassie-stained SDS-PAGE of supernatant and pellet fraction after Microtubule spin down. Using microtubule associated protein rich fraction (MAPF) as positive and BSA as negative control. Microtubule spin down assay of GST-His-CaM2 (I).

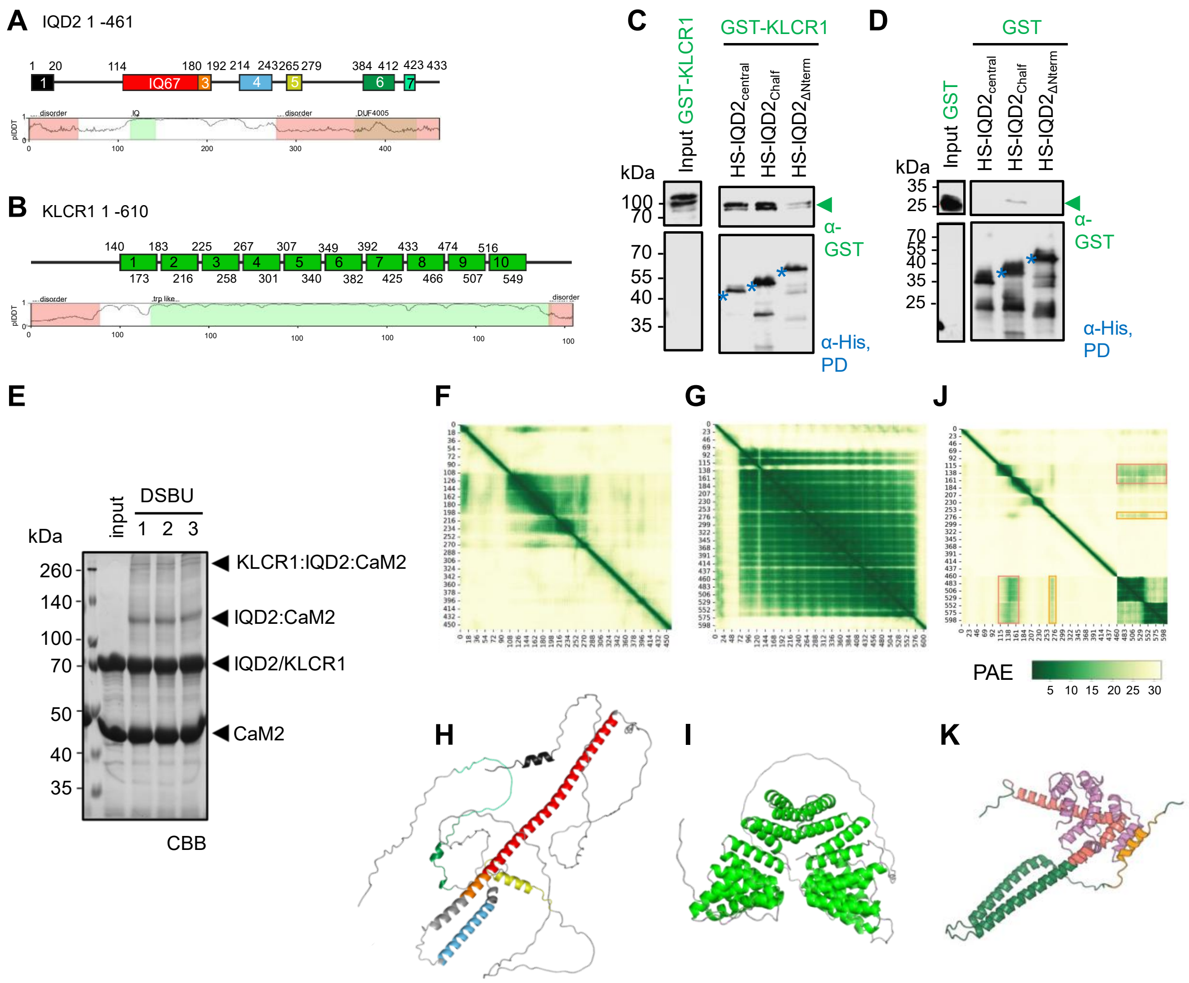


**Figure S3 (related to Figure 3): Structural predictions and structural determinants of IQD2-KLCR1 binding.**

**A,B,** Protein sequences of IQD2 (A) and of KLCR1 (B) with predicted motifs and domains shown as colored boxes (top). Known domains are highlighted in green (bottom). Disorder prediction is based on the MobiDB-lite consensus disorder prediction tool. The AlphaFold plDDT score is shown as line-plot inside the protein boxes. **C, D,** Pull-down assays with Ni-NTA-immobilized HS-IQD2 variants and GST-KLCR1 (C) and the free GST control (D). Western blot detection using anti-His and anti-GST antibodies. **E,** Preparative SDS-PAGE analysis of crosslinking reactions of KLCR1 and IQD2 with CaM2. Coomassie Brillant blue (CBB) stained gel of input containing HS-IQD2/CaM2/KLCR1 and three independent cross-linking (XL) reactions (1-3) after addition of DSBU. **F-I,** AlphaFold2 prediction of the monomeric IQD2 (F,H) and KLCR1 (G,I). High scores in PAE-plots (F,H) indicate a high confidence in spatial placement of the residue. Alphafold2 models of monomeric IQD2 (H) and KLCR1 (I) with motifs and domains color-coded according to A and B, respectively. **J,K**, CaM-modulated conformational change. PAE heatmap of IQD2:CaM2 Alphafold prediction. Binding regions stabilized are highlighted by colored boxes (J). AlphaFold2 model of IQD2 and CaM2 (K). Binding regions of higher confidence are colored salmon and orange.

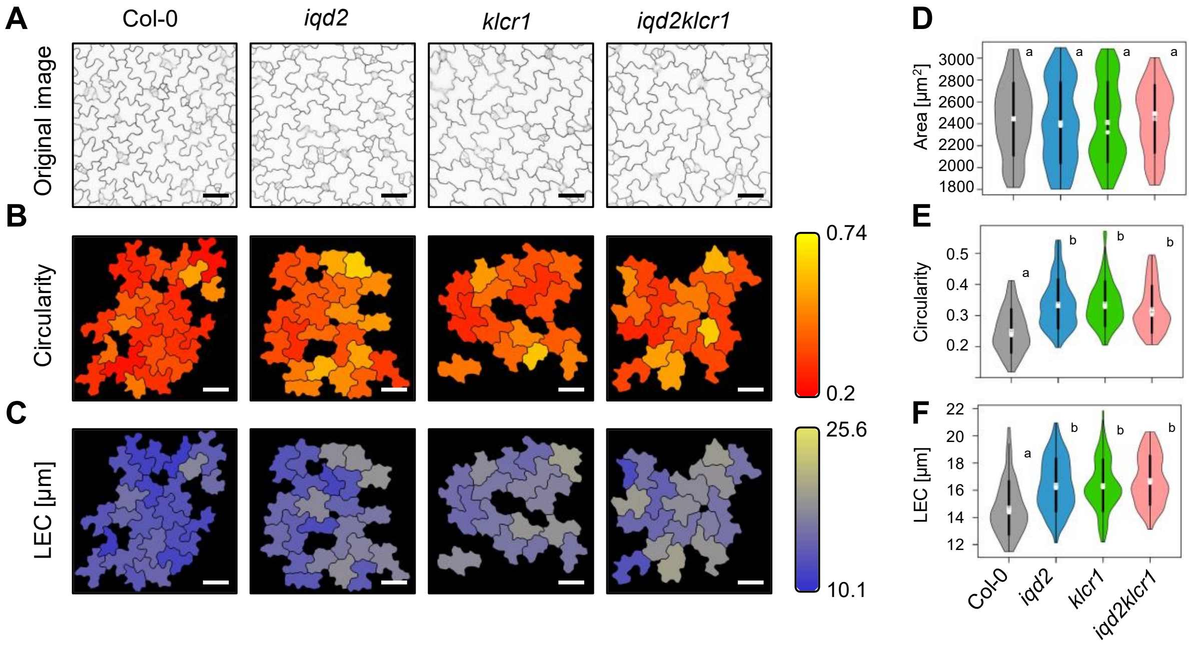


**Figure S4 (related to Figure 4): Pavement cell shape in wild type, *iqd2, klcr1* and *iqd2klcr1* mutants**

**A-C,** Pavement cells (PCs) in the abaxial side of cotyledons in 5-day-old seedlings of wild type, *iqd2, klcr1* and *iqd2klcr1* mutants. Images are single optical sections with cell contours labeled with PI (A). Heatmaps of circularity (B) and largest empty circle (LEC) (C). Scale bars, 50 µm. **D-F**, Quantitative analysis of PC features with PaCeQuant from n= 63 (Col-0), n=78 (*iqd2*), n=77 (*klcr1*), and n=60 (*iqd2klcr1*) cells, analyzed in six cotyledons from six seedlings. Violin plots of feature distributions of area (D), circularity (E), and LEC (F). Different letters denote significant statistical differences ($p < 0.001$; Kruskal-Wallis post-hoc Dunn‘s test).

**Table S1: Primers for cloning and genotyping:**

| Primer | # | Sequence (5' -> 3') |
|---|---|---|
| IQD2.1 fwd | 2461 | TTTGAATTCCTGGTGCCACGCGGTAGTATGGGGAAAAAACTAAAT |
| IQD2.1 rev | 2462 | TTTCTCGAGCTCTAAGGAGCAGATGAAGATG |
| IQD2.2 fwd | 2463 | TTTGAATTCCTGGTGCCACGCGGTAGTGGTGTTGTTCGTCG |
| IQD2.3 rev | 2466 | TTTCTCGAGCTCTAGGTCCCTCTTGCGG |
| IQD2.4 fwd | 2467 | TTTGAATTCCTGGTGCCACGCGGTAGTCCAAGAAACAAAAACAGTT T |
| IQD2.4 rev | 2468 | TTTTCTAGAT-TCAGCTGCCTGCT |
| IQD2.9 fwd | 2542 | GGGGACAAGTTTGTACAAAAAAGCAGGCTTCCTGGTGCCACGCGG TAGTGGTGTTGTTCGTCGCGCT |
| IQD2.3 new fwd | 2757 | GGGGACAAGTTTTACAAAAAAGCAGGCTTCATGTCAGAAGAGAAT CAGGCTCGC |
| IQD2.7 rev | 2759 | GGGGACCACTTTGTACAAGAAAGCTGGGTCTCAGCTGCCTGCTCC G |
| KLCR1 pET41a(+) fwd | 2523 | CGGTGATGACGACGACAAGAGTCCCATGGGAATGCCAGCAATGC CAGG |
| KLCR1 pET41a(+) rev | 2524 | GTGGTGGTGGTGCTCGAGTGCGGCCGCTCAGAACTTGAAACCGA GGC |
| CaM2 pET41a(+) fwd | 2525 | CGGTGATGACGACGACAAGAGTCCCATGGCGGATCAGCTCAC |
| CaM2 pET41a(+) rev | 2526 | GTGGTGGTGGTGCTCGAGTGCGGCCGCTCACTTAGCCATCATAAC CTTCA |
| Genotyping *klcr1* fwd | 286 | TATACAATAAGCCGGTGCCTG |
| Genotyping *klcr1* rev | 287 | GATCAATCAGATTCTGGAGCG |
| Genotyping *klcr2* fwd | 375 | GATTTGGAGTTGAGGCATTTG |
| Genotyping *klcr2* rev | 376 | CCAAAGCAACAGAGGAATGAG |
| Genotyping *iqd2* fwd | 73 | ATCCGAGCTAGGAGAATCAGG |
| Genotyping *iqd2* rev | 74 | GTTTAGCCTTGAGGGAGTTGG |
| SALK LB | A004 | ATTTTGCCGATTTCGGAAC |
| SAIL LB | A002 | GCCTTTTCAGAAATGGATAAATAGCCTTGCTTCC |
| GK LB | A009 | ATAATAACGCTGCGGACATCTACATTT |

**Table S2: Primer combinations for generation of expression constructs and genotyping**

| Protein | AA positions | Trivial name | Primers | Vector |
|---|---|---|---|---|
| KLCR1 | 1 to 610 | N/A | 2523 + 2524 | pET41a(+) |
| CaM2 | 1 to 149 | N/A | 2525 + 2526 | pET41a(+) |
| IQD2 | 1 to 461 | N/A | 2461 + 2468 | pENTR3C |
| | 1 to 97 | N-term | 2461 + 2462 | pENTR3C |
| | 328 to 461 | C-term | 2467 + 2468 | pENTR3C |
| | 98 to 327 | central | 2463 + 2466 | pENTR3C |
| | 98 to 461 | ΔN-term | 2542 + 2544 | pDONR221 |
| | 181 to 461 | C-half | 2757 + 2759 | pDONR221 |
| **Allele** | **Size (bp)** | | **Primers** | |
| *KLCR1* WT allele | 1113 | | 286 + 287 | |
| *klcr1* T-DNA | 503-703 | | 286 + A002 | |
| *KLCR2* WT allele | 1024 | | 375 + 376 | |
| *klcr2* T-DNA | 497-797 | | 375 + A004 | |
| *IQD2* WT allele | 928 | | 73 + 74 | |
| *iqd2* T-DNA | 406-606 | | 74 + A009 | |

#### References


1. Zang, J., Klemm, S., Pain, C., Duckney, P., Bao, Z., Stamm, G., Kriechbaumer, V., Bürstenbinder, K., Hussey, P.J. and Wang, P. (2021). A novel plant actin-microtubule

bridging complex regulates cytoskeletal and ER structure at ER-PM contact sites. Curr Biol *31*, 1251-1260.

2. Ganguly, A., Zhu, C., Chen, W. and Dixit, R. (2020). FRA1 kinesin modulates the lateral stability of cortical microtubules through Cellulose Synthase-Microtubule Uncoupling proteins. Plant Cell *32*, 2508–2524.
3. Gutierrez, R., Lindeboom, J.J., Paredez, A.R., Emons, A.M.C. and Ehrhardt, D.W. (2009). Arabidopsis cortical microtubules position cellulose synthase delivery to the plasma membrane and interact with cellulose synthase trafficking compartments. Nat Cell Biol *11*, 797–806.
4. Eng, R.C., Schneider, R., Matz, T.W., Carter, R., Ehrhardt, D.W., Jönsson, H., Nikoloski, Z. and Sampathkumar, A. (2021). KATANIN and CLASP function at different spatial scales to mediate microtubule response to mechanical stress in Arabidopsis cotyledons. Curr Biol *31*, 3262-3274.e6.
5. Bürstenbinder, K., Savchenko, T., Müller, J., Adamson, A.W., Stamm, G., Kwong, R., Zipp, B.J., Dinesh, D.C. and Abel, S. (2013). Arabidopsis Calmodulin-binding protein IQ67-Domain 1 localizes to microtubules and interacts with Kinesin Light Chain-related protein-1. J Biol Chem *288*, 1871–1882.
6. Bürstenbinder, K., Möller, B., Plötner, R., Stamm, G., Hause, G., Mitra, D. and Abel, S. (2017). The IQD family of Calmodulin-binding proteins links calcium signaling to microtubules, membrane subdomains, and the nucleus. Plant Physiol *173*, 1692–1708.
7. Fischer, C., Kugler, A., Hoth, S. and Dietrich, P. (2013). An IQ domain mediates the interaction with calmodulin in a plant cyclic nucleotide-gated channel. Plant Cell Physiol *54*, 573–584.
8. Karimi, M., Inzé, D. and Depicker, A. (2002). GATEWAY vectors for Agrobacterium-mediated plant transformation. Trends Plant Sci *7*, 193–195.
9. Nakagawa, T., Suzuki, T., Murata, S., Nakamura, S., Hino, T., Maeo, K., Tabata, R., Kawai, T., Tanaka, K., Niwa, Y., et al. (2007). Improved Gateway binary vectors: high-performance vectors for creation of fusion constructs in transgenic analysis of plants. Biosci Biotechnol Biochem *71*, 2095–2100.
10. Rajagopala, S.V., Hughes, K.T. and Uetz, P. (2009). Benchmarking yeast two-hybrid systems using the interactions of bacterial motility proteins. Proteomics *9*, 5296–5302.
11. Nguyen, A.N., Song, J.-A., Nguyen, M.T., Do, B.H., Kwon, G.G., Park, S.S., Yoo, J., Jang, J., Jin, J., Osborn, M.J., et al. (2017). Prokaryotic soluble expression and purification of bioactive human fibroblast growth factor 21 using maltose-binding protein. Sci Rep *7*, 16139.
12. Quan, J. and Tian, J. (2011). Circular polymerase extension cloning for high-throughput cloning of complex and combinatorial DNA libraries. Nat Protoc *6*, 242–251.
13. Kumari, P., Dahiya, P., Livanos, P., Zergiebel, L., Kölling, M., Poeschl, Y., Stamm, G., Hermann, A., Abel, S., Müller, S., et al. (2021). IQ67 DOMAIN proteins facilitate preprophase band formation and division-plane orientation. Nat Plants *7*, 739–747.
14. Ruhnow, F., Persson, S. and Schneider, R. (2023). Noninvasive long-term imaging of the cytoskeleton in Arabidopsis seedlings. Methods Mol Biol *2604*, 297–309.
15. Schindelin, J., Arganda-Carreras, I., Frise, E., Kaynig, V., Longair, M., Pietzsch, T., Preibisch, S., Rueden, C., Saalfeld, S., Schmid, B., et al. (2012). Fiji: an open-source platform for biological-image analysis. Nat Methods *9*, 676–682.
16. Möller, B., Poeschl, Y., Plötner, R. and Bürstenbinder, K. (2017). PaCeQuant: A tool for high-throughput quantification of pavement cell shape characteristics. Plant Physiol *175*, 998–1017.
17. Poeschl, Y., Möller, B., Müller, L. and Bürstenbinder, K. (2020). User-friendly assessment of pavement cell shape features with PaCeQuant: Novel functions and tools. Methods Cell Biol *160*, 349–363.

18. Mitra, D., Klemm, S., Kumari, P., Quegwer, J., Möller, B., Poeschl, Y., Pflug, P., Stamm, G., Abel, S. and Bürstenbinder, K. (2018). Microtubule-associated protein IQ67 DOMAIN5 regulates morphogenesis of leaf pavement cells in *Arabidopsis thaliana*. J Exp Bot *70*, 529–543.
19. Möller, B., Poeschl, Y., Klemm, S. and Bürstenbinder, K. (2019). Morphological analysis of leaf epidermis pavement cells with PaCeQuant. Methods Mol Biol *1992*, 329–349.
20. Möller, B., Glaß, M., Misiak, D. and Posch, S. (2016). MiToBo - A toolbox for image processing and analysis. J Open Res Softw *4*, e17.
21. Boudaoud, A., Burian, A., Borowska-Wykręt, D., Uyttewaal, M., Wrzalik, R., Kwiatkowska, D. and Hamant, O. (2014). FibrilTool, an ImageJ plug-in to quantify fibrillar structures in raw microscopy images. Nat Protoc *9*, 457–463.
22. Varapparambath, V., Mathew, M.M., Shanmukhan, A.P., Radhakrishnan, D., Kareem, A., Verma, S., Ramalho, J.J., Manoj, B., Vellandath, A.R., Aiyaz, M., et al. (2022). Mechanical conflict caused by a cell-wall-loosening enzyme activates de novo shoot regeneration. Dev Cell *57*, 2063-2080.
23. Kim, J., Harter, K. and Theologis, A. (1997). Protein-protein interactions among the Aux/IAA proteins. Proc Natl Acad Sci U S A *94*, 11786–11791.
24. Zhu, T., Sata, M. and Ikebe, M. (1996). Functional expression of mammalian myosin I beta: analysis of its motor activity. Biochem *35*, 513–522.
25. Iacobucci, C., Götze, M., Ihling, C.H., Piotrowski, C., Arlt, C., Schäfer, M., Hage, C., Schmidt, R. and Sinz, A. (2018). A cross-linking/mass spectrometry workflow based on MS-cleavable cross-linkers and the MeroX software for studying protein structures and protein-protein interactions. Nat Protoc *13*, 2864–2889.
26. Ihling, C.H., Piersimoni, L., Kipping, M. and Sinz, A. (2021). Cross-linking/mass spectrometry combined with ion mobility on a timsTOF pro instrument for structural proteomics. Anal Chem *93*, 11442–11450.
27. Götze, M., Pettelkau, J., Fritzsche, R., Ihling, C.H., Schäfer, M. and Sinz, A. (2015). Automated assignment of MS/MS cleavable cross-links in protein 3D-structure analysis. J Am Soc Mass Spectr *26*, 83–97.
